# Activity-regulated gene expression across cell types of the mouse hippocampus

**DOI:** 10.1101/2022.11.23.517593

**Authors:** Erik D. Nelson, Kristen R. Maynard, Kyndall R. Nicholas, Matthew N. Tran, Heena R. Divecha, Leonardo Collado-Torres, Stephanie C. Hicks, Keri Martinowich

**Author notes:** equal contribution (Stephanie C. Hicks, Keri Martinowich). **Correspondence**, Stephanie C. Hicks, Keri Martinowich.

## Abstract

Activity-regulated gene (ARG) expression patterns in the hippocampus (HPC) regulate synaptic plasticity, learning, and memory, and are linked to both risk and treatment response for many neuropsychiatric disorders. The HPC contains discrete classes of neurons with specialized functions, but cell type-specific activity-regulated transcriptional programs are not well characterized. Here, we used single-nucleus RNA-sequencing (snRNA-seq) in a mouse model of acute electroconvulsive seizures (ECS) to identify cell type-specific molecular signatures associated with induced activity in HPC neurons. We used unsupervised clustering and *a priori* marker genes to computationally annotate 15,990 high-quality HPC neuronal nuclei from *N*=4 mice across all major HPC subregions and neuron types. Activity-induced transcriptomic responses were divergent across neuron populations, with dentate granule cells being particularly responsive to activity. Differential expression analysis identified both upregulated and downregulated cell type-specific gene sets in neurons following ECS. Within these gene sets, we identified enrichment of pathways associated with varying biological processes such as synapse organization, cellular signaling, and transcriptional regulation. Finally, we used matrix factorization to reveal continuous gene expression patterns differentially associated with cell type, ECS, and biological processes. This work provides a rich resource for interrogating activity-regulated transcriptional responses in HPC neurons at single-nuclei resolution in the context of ECS, which can provide biological insight into the roles of defined neuronal subtypes in HPC function.

## 1 Introduction

Learning and memory are associated with changes in neuronal firing patterns that occur in response to experience. These physiological processes are modulated at the cellular and molecular levels through alterations in dendritic morphology, receptor trafficking, neurotransmitter release dynamics, and synaptic plasticity (Bailey et al., 2015; Diering and Huganir, 2018; Palacios-Filardo and Mellor, 2019). Activity-regulated genes (ARGs) are rapidly and transiently transcribed in response to neural activity and critically mediate many cellular modifications that control these processes (Gallo et al., 2018; Gray and Spiegel, 2019; Tyssowski and Gray, 2019). Transcription of these genes occurs in multiple waves, with rapid primary response genes (rPRGs) and delayed primary response genes (dPRGs) preceding secondary response genes (SRGs), although timing can vary according to activity duration (Tyssowski et al., 2018). Crucially, recent single-cell RNA-sequencing (scRNA-seq) and single-nucleus RNA-sequencing (snRNA-seq) studies in mouse amygdala and cortex, respectively, have found cell type-specific upregulation of ARGs following induction of electroconvulsive seizures (ECS) (Hu et al., 2017; Wu et al., 2017).

In the hippocampus (HPC), ARG expression mediates robust changes in synaptic plasticity, which controls multiple forms of hippocampal-dependent learning and memory, and is implicated in neuropsychiatric disorders as well as their treatment response. For example, ARG induction in HPC neurons is hypothesized to be a key molecular mechanism underlying efficacy of antidepressant treatments, including electroconvulsive therapy (ECT) (Castrén and Monteggia, 2021; Chottekalapanda et al., 2020; Nagy et al., 2018). Several studies have examined ARG expression in HPC following acute ECS induction (Newton et al., 2003; Kodama et al., 2005;Aoyama et al., 2022; Altar et al., 2004; Newton et al., 2003; Ploski et al., 2006; Kodama et al., 2005). However, these studies either lack single-cell resolution or consider only targeted sets of genes. Multiple studies have meticulously characterized gene expression of cell types in mouse HPC (Yao et al., 2021; Habib et al., 2016;Habib et al., 2017; Zeisel et al., 2015; Ding et al., 2020; Cembrowski et al., 2016b; Cembrowski et al., 2018), yet cell type-specificity of gene expression changes in response to neural activity has not been investigated at transcriptome-wide scale within HPC. This is important because specialized cell types and subregions within the HPC are linked to specific molecular, cellular, circuit, and behavioral-level functions, and are selectively vulnerable in different pathological conditions (Pelkey et al., 2017; Ayhan et al., 2021; Alkadhi, 2019; Dengler and Coulter, 2016; Collado-Torres et al., 2019).

In this study, we investigated cell type-specific molecular expression patterns in mouse HPC neurons following induction of neural activity after ECS. Specifically, we used snRNA-seq to assess transcriptome-wide gene expression in HPC neuronal cell types under both Sham and ECS conditions at cellular resolution. We performed both sample-level and cell type-specific pseudobulk differential expression analysis. We found particularly strong differential expression between Sham and ECS in dentate granule cells (GCs), and identified heterogeneous activity-regulated gene expression signatures across HPC neurons. Using non-negative matrix factorization, we also found continuous patterns of gene expression associated with ECS across HPC neurons. We provide an interactive web application for exploration of this data to further advance understanding of the heterogeneity of activity-regulated gene expression in the HPC.

## 2 Materials and Methods

### 2.1 Animals

C57Bl6/J mice (male, stock #000664, Jackson Laboratories) were group-housed (3-5 animals per cage) and maintained on a 12 hour light/dark cycle in a temperature and humidity-controlled colony room. All animals had *ad libitum* access to food and water. Animals were male, 10-16 weeks of age, and weighed 25-35 g at study initiation. All procedures were in accordance with the Institutional Animal Care and Use Committee of the Johns Hopkins University School of Medicine.

### 2.2 Electroconvulsive seizures (ECS) and mouse brain tissue collection

Mice were administered either Sham or electroconvulsive seizures (ECS) as previously described (Maynard et al., 2018). Mice were anesthetized using inhaled isoflurane prior to and during Sham and ECS treatment. ECS were delivered with an Ugo Basile pulse generator (model #57800-001, shock parameters: 100 pulse/s frequency, 3 ms pulse width, 1 s shock duration and 50 mA current) (Maynard et al., 2018; Schloesser et al.,2015). Seizures were confirmed by visualization of tonic-clonic convulsion. Ninety minutes after treatment, animals were killed, brains were removed from the skull, and the HPC was dissected out of the brain on wet ice. Hippocampal tissue was flash-frozen in 2-methylbutane (ThermoFisher), and stored at −80°C until processing for snRNA-seq.

### 2.3 snRNA-seq data generation

We performed single-nucleus RNA-sequencing (snRNA-seq) on 4 samples using the 10× Genomics Chromium Single Cell Gene Expression V3 technology (Zheng et al., 2017). Nuclei were isolated using the “Frankenstein” nuclei isolation protocol developed for frozen tissues, as described in the customer-developed protocol from 10× Genomics (Habib et al., 2016; Habib et al., 2017; Hu et al., 2017; Lacar et al., 2016; Tran et al., 2021). Briefly, ~40mg of frozen HPC tissue from each sample was homogenized in chilled Nuclei EZ Lysis Buffer (MilliporeSigma #NUC101) using a glass dounce with ~10-15 strokes per pestle. Homogenates were filtered using 70 μm mesh strainers and centrifuged at 500 × *g* for 5 minutes at 4°C in a benchtop centrifuge. Nuclei were resuspended in EZ lysis buffer, centrifuged, and equilibrated in nuclei wash/resuspension buffer (1x PBS, 1% BSA, 0.2U/μL RNase Inhibitor). Nuclei were rinsed and centrifuged in the nuclei wash/resuspension buffer 3 times. Next, they were labeled with propidium iodide (PI) and Alexa Fluor 488-conjugated anti-NeuN (MilliporeSigma cat. #MAB377X), at 1:1000 in the same wash/resuspension buffer for 30 minutes on ice. NeuN-labeling was used to facilitate enrichment of neurons during fluorescent-activated nuclear sorting (FANS). Following labeling, samples were filtered through a 35 μm cell strainer and sorted on a BD FACS Aria II Flow Cytometer (Becton Dickinson) into 10X Genomics reverse transcription reagents. Gating criteria were hierarchically selected for whole, singlet nuclei (by forward/side scatter), G0/G1 nuclei (by PI fluorescence), and NeuN-positive cells. For each sample, approximately 5,000 single nuclei were sorted directly into 25.1 μL of reverse transcription reagents from the 10× Genomics Single Cell 3’ Reagents kit (without enzyme). The 10× Chromium process was performed and libraries were prepared according to manufacturer’s instructions for the V3 3’ gene expression kit (10× Genomics). Samples were sequenced on a Next-seq (Illumina) at the Johns Hopkins University Single Cell and Transcriptomics Sequencing Core.

### 2.4 snRNA-seq preprocessing, quality control, and doublet removal

Using the raw FASTQ files, we used 10× Genomics’ `cellranger count` version 3.0.2 (https://support.10xgenomics.com/single-cell-gene-expression/software/release-notes/3-0) to map the reads to the mouse reference transcriptome mm10 version 3.0.0. Alignment was performed using a modified “pre - mRNA” GTF in order to include both intronic and exonic reads. (https://support.10xgenomics.com/single-cell-gene-expression/software/pipelines/latest/advanced/references#premrna). For analysis steps, we used version 4.1.3 of R and version 3.14 of the Bioconductor suite of R packages for analysis of genomics data (Huber et al., 2015, Amezquita et al., 2020). We read in the nucleus × gene UMI count matrices as a `singleCellExperiment` object using the `read10xCounts` function from the `DropletUtils` package and `singleCellExperiment` (Amezquita et al., 2020) packages.

For quality control, we first used the `emptyDrops` function from the `DropletUtils` package (Lun et al.,2019), taking a Monte Carlo simulation-based approach to identify empty droplets and other sources of ambient RNA, such as debris. For the remaining steps, we then used the `scater` and `scran` packages for thresholding nuclei by mitochondrial expression rate, library size, and number of detected genes (McCarthy et al., 2017; Lun et al., 2016a; Lun et al., 2019). More specifically, we used the `perCellMetrics` function from scran to compute per-cell quality control metrics, and with the `isOutlier` function from `scater`, we applied a 3× median absolute deviation (MAD) threshold to remove nuclei with a high percent of UMIs mapping to mitochondrial genes, low library sizes, or a small number of expressed genes. Additionally, we computed doublet scores for all nuclei per sample,using the `computeDoubletDensity` function from the `scDblFinder` package (Germain et al., 2021), discarding those nuclei with a doublet score greater than 5. This discarded approximately 1.8% of the nuclei, slightly more conservative than 10× Genomics estimate of approximately 2-3% based on the number of nuclei loaded per sample (10× Chromium Single Cell 3′ Reagent Kits User Guide v3.1). We also discarded genes with zero counts across all nuclei. After applying these QC steps, the number of genes × the number of nuclei was 25,562 × 17,804. Finally, with the described unsupervised clustering below, we removed thalamic cells to obtain a final matrix of 15,990 nuclei for downstream analyses (see Sections 2.7.2-3).

### 2.5 Feature selection and dimensionality reduction

For feature selection, we used the `scry` R/Bioconductor package (Townes et al., 2019) to identify the top 3,000 highly deviant genes (HDGs) using a Poisson model fit to the gene expression for each gene across all cells.

For dimensionality reduction, we fit a Poisson model to gene counts and calculate Pearson residuals for each gene. Principal component analysis (PCA) was then applied to the Pearson residuals. This approach approximates GLM-PCA, a feature selection and dimensionality reduction method shown to operate in a more statistically sound manner to the nature of droplet-UMI snRNA-seq data than other widely used methods (Townes et al., 2019). We retained 50 principal components (PCs) for downstream analyses.

For the purpose of visualization, we used the implementation of uniform manifold approximation and projection (UMAP) (McInnes et al., 2018) in `scater` to generate a low-dimensional visual representation of the high-dimensional data. We combined this method with the `schex` package in order to mitigate visualization difficulties caused by overplotting (Freytag and Lister, 2020). This package allows for binning of data into hexagon-shaped bins within reduced dimension plots, thereby allowing for better visualization of sample-level metadata variables, such as condition or cell-type annotation.

### 2.6 Normalization

We used the scran package for normalization of expression values across nuclei. Specifically, we first used the `quickCluster` function to cluster similar cells based on expression profiles. Then, we computed size factors from these preliminary clusters using the `computeSumFactors` function. This function sums all counts for each cluster and computes a library size factor by dividing this summed profile by the average of counts across all cells. Lastly, we used the computed size factors and the `logNormCounts` function to normalize counts and apply a log_2_ transformation (Lun et al., 2016a).We used these normalized counts for visualization purposes only, and not for feature selection, or dimensionality reduction, or clustering (Pearson residuals-based).

### 2.7 Clustering

For clustering, we applied graph-based clustering to the top 50 PCs. More specifically, we first used the `buildSNNGraph` function from scran to build a shared-nearest neighbors graph with *k*=50 nearest neighbors. This function is a wrapper for the `makeSNNGraph` function from the `bluster` package (Lun, 2022), which identifies *k* nearest neighbors between nuclei based on Euclidean distances between their gene expression profiles. Edges are drawn between nuclei with at least one shared-nearest neighbor using a rank-based weighting method (Xu and Su, 2015). Following graph construction, we used the `cluster_walktrap` method from CRAN package `igraph` (Csardi and Nepusz, 2005) for random walk-based community detection within our graph. This approach yielded 21 preliminary clusters.

#### 2.7.1 Cluster marker gene detection

We identified marker genes for each preliminary cluster with the `findMarkers` function from `scran`. More specifically, we used `findMarkers` to perform pairwise *t*-tests between all pairs of clusters. We set arguments `direction=“up”` and `pval.type=“all”`, thereby only considering genes with a positive log fold change and combining *p*-values across all pairwise comparisons for a given cluster. This allowed us to generate lists of specific marker genes upregulated in one cluster when compared with all other clusters.

#### 2.7.2 Thalamic cluster identification and removal

After analyzing marker genes, we found genes known to be enriched in thalamus or epithalamus in marker gene lists for three preliminary clusters: cluster 5 (*Six3, Sox14, Zfhx3*), cluster 9 (*Shox2, Rorb, Zfhx3*), and cluster 14 (*Ano1, Chat, Pou4f1*) (Nagalski et al., 2016). Additionally, each of these clusters also expressed *Tcf7l2*, a transcription factor highly specific to thalamic and epithalamic nuclei (Lee et al., 2017). As our study is focused on HPC, these nuclei were not included in downstream analyses and we removed all nuclei in preliminary clusters 5, 9, and 14 from the *SingleCellExperiment* object. However, the dataset that is publicly released on GitHub does include these nuclei.

#### 2.7.3 Reclustering

Following removal of thalamic clusters, the dimensions of our data set were 25,562 genes × 15,990 nuclei. This included 3,755 and 4,460 nuclei from the two Sham samples and 3,777 and 3,998 from the two ECS samples. We repeated the following steps as detailed above for these nuclei: feature selection, dimensionality reduction, normalization, UMAP embedding generation, clustering, and cluster marker identification. However, here we used *k*=25 for graph-based clustering to allow for higher resolution within HPC cell types. This yielded 28 clusters. Of these, using known marker genes for hippocampal regions – dentate gyrus, cornu ammonis (CA) subfields 1-4 and the more proximal retrohippocampal (RHP) regions such as subiculum and prosubiculum, we annotated 2 dentate granule cell clusters (*Prox1+*), 1 CA4 neuron cluster (*Calb2+*), 2 CA3 pyramidal cell clusters (*Nectin3+, Iyd+* for CA3.1, likely more superficial; *St18+* for CA3.2, likely deeper), 1 CA2 pyramidal cell cluster (*Ntsr2*+, *Rgs14+),* 1 CA1 pyramidal cell cluster (*Man1a+, Zdhhc2+*), 2 clusters corresponding with prosubiculum (PS) neurons (*Ntng2+, Nos1+, Dcn+* for PS.1; *Ntng2+, Ndst4+, Cntn6+, Fras1+* for PS.2) and 1 cluster corresponding with subiculum neurons (*Ndst4+, Fn1+, Nts+*) (Dong et al., 2009; Cembrowski et al., 2016b; Cembrowski et al., 2016a; Cembrowski et al., 2018; Habib et al., 2016; Bienkowski et al., 2018; Ding et al., 2020; Yao et al., 2021; Kalpachidou et al., 2021). All excitatory clusters from dentate gyrus and CA regions expressed high levels of transcription factor *Zbtb20* (Rosenthal et al., 2012). PS.1 also contains cells expressing *Gpc3*, suggesting it may contain a distinct population of neurons from the ventral tip of CA1 as well. These neurons are similar to PS neurons in both spatial proximity and gene expression (Ding et al., 2020; Yao et al., 2021). We annotated 5 clusters corresponding with different GABAergic neuron types: CGE-derived GABAergic neurons (*Htr3a+),* CGE-derived neurogliaform neurons (*Ndnf+, Lamp5+),* MGE-derived neurogliaform neurons (*Lhx6+, Lamp5+),* MGE-derived fast-spiking GABAergic neurons (*Pvalb+),* and MGE-derived *Sst*+ GABAergic neurons (Harris et al., 2018; Yao et al., 2021). The remaining 13 clusters corresponded with neurons of more distal RHP regions, such as presubiculum, postsubiculum, parasubiculum, area prostriata, and entorhinal cortex (Yao et al., 2021). To distinguish between these clusters, we observed expression of the following marker genes: *Cux1, Cux2, Satb2* for layers 2 and 3; *Pamr1* for layer 5/polymorphic (Po) layer; *Foxp2, Cobll1, Ctgf* for layers 6/6b (Ding et al., 2020; Yao et al., 2021). We grouped these 13 RHP clusters into the following categories: L2/3 (7 clusters), L5/Po (2 clusters), and L6/6b (4 clusters). Thus, for downstream analyses, we used 18 cell type groups or clusters, which we refer to as annotated cell type in subsequent methods sections. We also used the `iSEE` Bioconductor package to generate an interactive app for visualization of our dataset (Rue-Albrecht et al., 2018).

### 2.8 Differential expression analysis

#### 2.8.1 Differential expression analysis between sham and ECS

To perform differential expression (DE) analysis, we began by aggregating all the UMI counts for each genes across all nuclei within each sample (or mouse). This step is referred to as a “pseudobulk” sample. We pseudobulked the `SingleCellExperiment` object by sample using the `aggregateAcrossCells` function from scran, yielding a separate `SingleCellExperiment` object with dimensions of 25,562 genes × 4 samples. Next, we performed DE analysis using a pipeline constructed using the `edgeR` package (McCarthy et al., 2012; Robinson et al., 2010; Chen et al., 2016). We converted the pseudobulked `SingleCellExperiment` object into a `DGEList` object. We filtered lowly expressed genes with the `filterByExpr` function and normalized the pseudobulked data using the `calcNormFactors` function. We built the design matrix using only the condition (Sham and ECS) by calling `model.matrix(~condition`). We passed this design matrix and the `DGElist` object to the `estimateDisp` function to estimate common and trended negative binomial dispersions. Using `glmQLFit`, we also estimate quasi-likelihood (QL) dispersions and fit a negative binomial generalized log-linear model to the pseudobulked data (Lund et al.,2012). We then test for DE between conditions using the `glmQLFtest` function, which uses a QL *F*-test for better type 1 error rate detection, compared with Chi-square approximation of likelihood ratio (Lun et al.,2016b). We used the Benjamini-Hochberg false discovery rate (FDR) to correct *p*-values for multiple testing and use an FDR threshold of 0.05 for determining DE genes (Benjamini and Hochberg, 1995). We generated a table of statistics from this analysis for all genes using the `topTags` function.

#### 2.8.2 Cell-type specific differential expression analysis

For cell type-specific differential expression analysis, we again pseudobulked the `SingleCellExperiment` using `aggregateAcrossCells`, but this time, we grouped by both sample and annotated cell type. We then used the `pseudoBulkDGE` function from `scran` to loop the same `edgeR`-based DE analysis detailed above across each annotated cell type label. This returned a list of tables containing gene-level DE statistics for each cell type, as was done above with `topTags`. Next, we applied the same 0.05 FDR threshold to identify DE genes for each cell type grouping. Finally, we retained the list of gene-level statistics for use in downstream functional enrichment analysis.

### 2.9 Matrix Factorization

As an orthogonal approach to graph-based clustering to examine the impact of acute ECS on HPC neurons, we used non-negative matrix factorization (NMF) to identify continuous gene expression patterns within our data (Stein-O’Brien et al., 2019). The `CoGAPS` package (Sherman et al., 2020; Fertig et al., 2010) uses a Bayesian non-negative matrix factorization (NMF) algorithm to decompose a gene count matrix into amplitude (A) and pattern (P) matrices, which correspond with sources of variation across genes and samples, respectively. In order to reduce memory, we first removed genes in the counts matrix with a sum of less than 20 UMI counts across all nuclei. This resulted in a matrix with dimensions of 19,829 genes × 15,990 nuclei. We passed this matrix to the `CoGAPS` function with the following parameters: `sparseOptimization=T, nIterations=12000, nPatterns=70, nSets=20`. We parallelized the algorithm across 20 cores for better performance. This yielded a `cogapsResult` object with 57 patterns matched across sets where patterns correspond to variation across cells. For marker gene detection across patterns, we used the `patternMarkers` function from `CoGAPS` (Stein-O’Brien et al., 2017).

To explore the relationship between discovered patterns from CoGAPS and our annotated cell types, we used a similar strategy to that used by Stein-O’Brien et al, 2019 (Stein-O’Brien et al., 2019). First, we created a new one-hot encoded matrix from our `singleCellExperiment` object, one column for each of our annotated cell types. This matrix had dimensions of 15,990 nuclei × 18 annotated cell types, where 1 indicated that a given nucleus is annotated as a given cell type and 0 indicated that it is not annotated as the given cell type. We then found the correlation between this matrix and the pattern matrix from `CoGAPS`, which was obtained by calling the `sampleFactors` slot from the `CogapsResult` object. To explore the relationship between discovered patterns and condition, we performed the same procedure detailed above, using instead a one-hot encoded matrix with dimensions of 15,990 nuclei × 2 condition levels (Sham or ECS).

### 2.10 Functional enrichment analysis

#### 2.10.1 Over-representation analysis (ORA)

We used over-representation analysis (ORA) to determine functional enrichment of Gene Ontology (GO) gene sets within identified upregulated DE genes between conditions in our sample-level DE analysis (described above in Section 2.8). We used the `enrichGO` function from the `clusterprofiler` (Wu et al., 2021; Yu et al., 2012) Bioconductor package, where we input a vector of DE genes with the following parameters: `pAdjustMethod = “BH”`, `pvalueCutoff = 0.01, qvalueCutoff = 0.05`. That is, we used an FDR cutoff of 0.05 to assess the statistical significance of enriched gene sets. We applied this method to all biological process (BP) gene sets within the GO resource (Ashburner et al., 2000; The Gene Ontology Consortium, 2021).

#### 2.10.2 Gene set enrichment analysis (GSEA)

We used gene-set enrichment analysis (GSEA) (Subramanian et al., 2005) to determine enrichment of GO gene sets within cell type-specific DE analysis results. In contrast to using a vector of DE genes as input, GSEA takes a ranked vector of gene-level statistics, such as *t*-statistics or *Z*-scores, for all genes. This allows for more nuanced calculation of gene set enrichment, which was desired for this analysis.

For GSEA of cell type-specific DE results, we first created a named vector of genes for each cell type grouping, where each name is a gene and each value is a *t*-statistic for that gene, calculated for each gene using `sign(logFC) × sqrt(F)`, where `F` refers to the *F*-statistic for each gene. *F*-statistics are provided in the output from `pseudoBulkDGE`. Each vector is ranked by *t*-statistic prior to GSEA. We passed this list of ranked, named vectors to `clusterProfiler`’s `gseGO` function for the analysis using the following parameters: `minGSSize=100, maxGSSize= 500, pvalueCutoff=0.05, eps= 0`. We again use an FDR-adjusted *p*-value cutoff of 0.05 to assess the statistical significance of enriched patterns. We performed this analysis for all cell types for GO terms within BP, CC, and MF domains.

For all balloon plots involving GSEA results, we sized circles by -log_10_-transformed FDR-adjusted *p*-values and sized by gene ratio size. Gene ratio is calculated by dividing leading edge gene subset size by total gene set size for each term. Leading edge is also known as the core enrichment set of genes and refers to genes that contribute most to the enrichment result for a given gene set.

#### 2.10.3 Semantic similarity clustering of GSEA results

Because of the large number of GO terms discovered, we used the `rrvgo` package to cluster enriched GO terms from GSEA (Yu et al., 2010; Yu, 2020; Sayols, 2020). This method uses semantic similarity analysis to reduce redundancy of terms and allow for better interpretation of results. First, for each of the three GO domains, we pooled all terms found to be enriched in at least one cell type. We then used `rrvgo`’s `calculateSimMatrix` function to calculate similarity matrices for each list of terms using the Wang method of semantic similarity measurement (Wang et al., 2007). Next, we clustered terms using the `reduceSimMatrix` function using a similarity threshold of 0.9 for BP and 0.7 for CC and MF, where the threshold levels can range from 0 to 1 and higher values indicate higher levels of similarity between terms. Names for each cluster were assigned as the term within each cluster with the largest mean -log_10_ adjusted *p*-value across all cell types. Lower threshold levels were used for CC and MF to account for lower total number of statistically significant terms for these domains. We ranked terms within each cluster based on mean negative log-transformed, FDR-adjusted *p*-value for each term.

## 3 Results

### 3.1 Characterization of neurons in the mouse hippocampus

To characterize cell type-specific ARG expression in mouse HPC, we used a well-defined mouse model of electroconvulsive seizures (ECS) to induce robust neural activity in hippocampal neurons (Schloesser et al.,2015; Maynard et al., 2018). Mice were administered either Sham or ECS (*N*=2 mice per group), and tissue was collected and processed for snRNA-seq, 90 min following seizure induction (**Figure 1A**). We generated a total of 17,804 quality-controlled, snRNA-seq profiles across all 4 mouse samples, which we then analyzed in a cell type and condition-specific manner.

**Figure 1:**
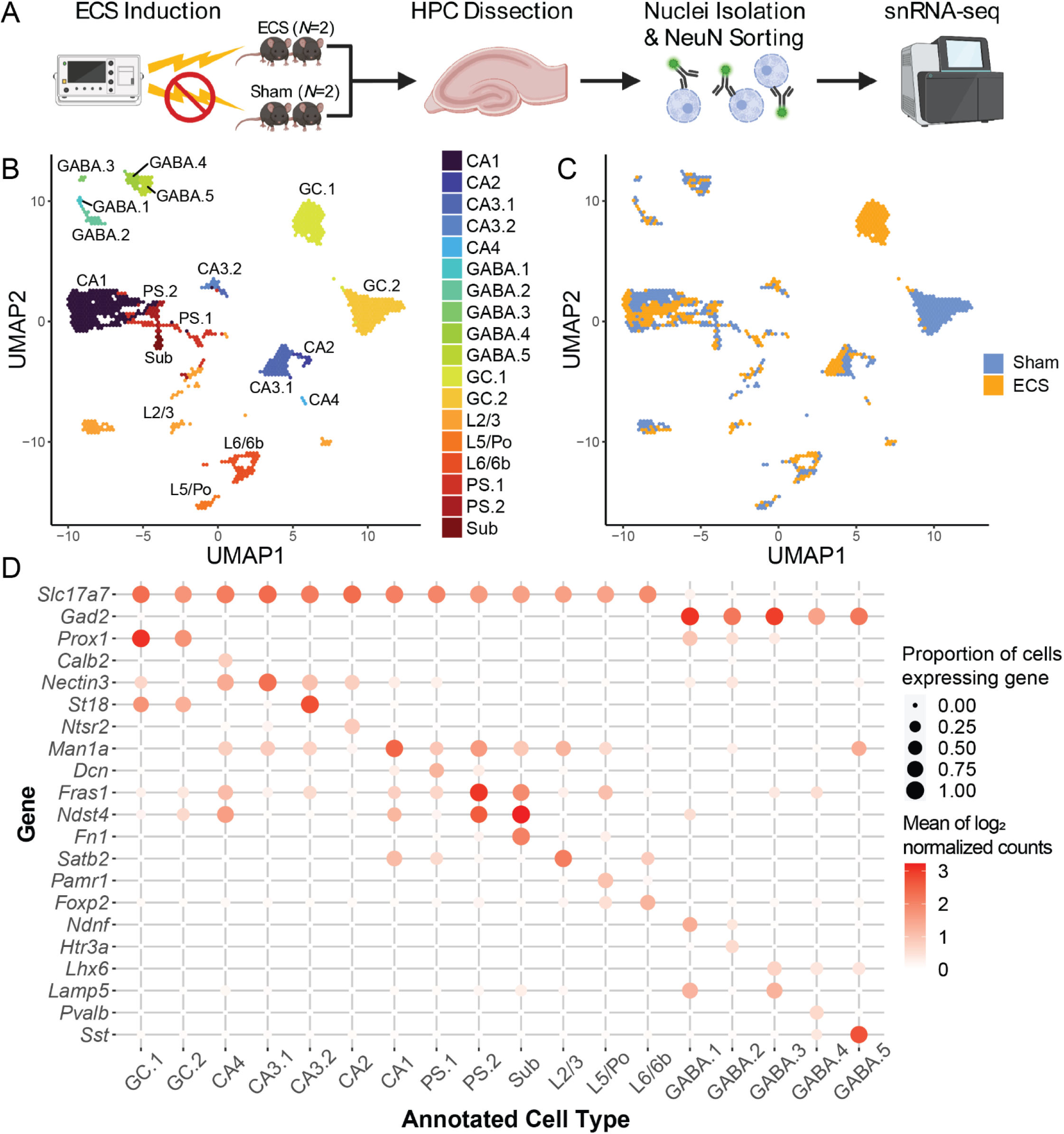
Distinct molecularly-defined cellular sub-populations in the mouse hippocampus. **(A)** Schematic of the experimental design to generate pre-quality control 17,804 nuclei from *N*=4 mice: *N*=2 Sham and *N*=2 electroconvulsive seizures (ECS). **(B)** UMAP representation of post-quality control 15,990 nuclei from the hippocampus (HPC) and retrohippocampal area (RHP) with color representing broad, related groups of cell types. 18 clusters/cluster groupings were identified using unsupervised graph-based clustering, and are plotted as annotated cell types. **(C)** UMAP representation with color representing experimental conditions. **(D)** Heatmap along a grid with circles colored by mean log_2_ expression of canonical gene markers averaged across nuclei (rows) for 18 clusters (columns). Circles are sized by the proportion of nuclei with nonzero expression values for each gene.

We applied standard preprocessing and quality control (QC) metrics including (i) removing doublets and (ii) removing nuclei with small library sizes, small numbers of expressed genes, or large numbers of UMIs mapping to mitochondrial genes (Section 2.4). Feature selection and dimensionality reduction were performed (Section 2.5), followed by an initial round of unsupervised clustering (Section 2.7) using nuclei from both Sham and ECS conditions to identify cell types. Graph-based clustering identified a total of 21 clusters. As expected due to neuronal enrichment during sorting, all clusters were of neuronal origin based on expression of neuron-specific markers, such as *Snap25* (**Figure S1A**). Of these clusters, we identified 3 as thalamic or epithalamic clusters based on expression of region-specific markers, most notably *Tcf7l2* and *Zfhx3,* which are thalamus and epithalamus-specific transcription factors (Nagalski et al., 2016) (**Figure S1B-F**). These clusters contained a total of 1,804 nuclei and were excluded from downstream analysis.

After removing thalamic clusters, we retained 15,590 high-quality nuclei. We performed a second round of feature selection, dimensionality reduction, and graph-based clustering to identify 28 distinct clusters, which were then merged into 18 groups (**Figure 1B**) as described in Section 2.7. Nearly all clusters contained nuclei from Sham and ECS conditions, with the exception of the dentate granule cell (GC) clusters. GC.1 was primarily composed of nuclei from ECS animals while GC.2 was primarily composed of nuclei from Sham animals (**Figure 1C**). We identified marker genes for all clusters (**Table S1**). We identified glutamatergic clusters mapping to known HPC neuron types, including pyramidal neurons from cornu ammonis (CA) subfields, CA4/hilar mossy cells, and GCs. Additionally, 3 clusters were mapped to more proximal RHP subregions, with 2 mapping to prosubiculum (PS) and 1 to subiculum (Sub). The remaining 13 glutamatergic clusters corresponded with more distal RHP subtypes, such as presubiculum, postsubiculum, and entorhinal cortex. These clusters were grouped based on assignment to L2/3 (7 clusters), L5/Po (2 clusters), and L6/6b (4 clusters). The remaining 5 clusters expressed *Gad2* and were annotated as marker-specific GABAergic subtypes (Section 2.7) (**Figure 1D**). GC and CA clusters are distinct in expression of transcription factor *Zbtb20* (**Figure S2A**), while PS and Sub clusters could be distinguished based on expression of known marker genes (**Figure S2B-H**). Clusters from RHP L2/3, L5/Po, and L6/6b were grouped based on shared marker gene expression (**Figure S2I-P**). This analysis demonstrates that our dataset comprises all neuronal cell types expected to reside across the mouse HPC.

### 3.2 Induction of robust activity-regulated gene expression in the hippocampus following electroconvulsive seizures

To identify activity-regulated genes (ARGs), we performed differential expression (DE) analysis between the Sham and ECS conditions. We first pseudobulked the data by summing UMI counts for each gene across all nuclei within each sample (*N*=2 Sham, *N*=2 ECS). This approach allowed us to make use of well-characterized bulk RNA-seq DE methods (McCarthy et al., 2012). Following pseudo-bulking, we used the *edgeR* Bioconductor package to perform DE analysis (Robinson et al., 2010). Although this initial analysis did not consider cell type, it increased statistical power by including all cells in one analysis. Additionally, it provided a broad picture of differential expression, which is useful for comparison to our cell type-specific analyses.

This analysis identified 789 genes at an FDR threshold of 0.05. Of these genes, 243 had a log_2_ fold change < 0 (downregulated) while 546 had a log_2_ fold change > 0 (upregulated) (**Figures 2A & S3; Table S2**). We compared these differentially expressed genes (DEGs) to a curated list of known ARGs (Tyssowski et al.,2018), and found 55 of these ARGs to be differentially expressed following ECS. Genes from all 3 classes of ARGs were upregulated: 6 rPRGs (including *Arc, Egr3, Gadd45g*, and *Junb*), 38 dPRGs (such as *Baz1a, Brinp1, Grasp*, and *Bdnf*), and 9 SRGs (such as *Nptx2, Mfap3l, Kcna1, Kcnj4*) (**Figure 2B**). We next performed gene ontology (GO) over-representation analysis to investigate enrichment of specific biological processes associated with upregulated DEGs (Yu et al., 2012). This revealed a total of 484 terms at an FDR threshold of 0.05 (**Table S3**). Top enriched GO terms for upregulated DE genes included growth, synapse, and plasticity associated processes such as synapse organization, synaptic vesicle cycle, and neurotransmitter secretion (**Figure 2C**). Additionally, we found enrichment of terms associated with transforming growth factor beta signaling, indicating potential involvement of immune system pathways in ECS. Overall, these data illustrate robust induction of activity-dependent gene expression by ECS across multiple classes of ARGs that implicate broad categories of synaptic function in mouse HPC.

**Figure 2:**
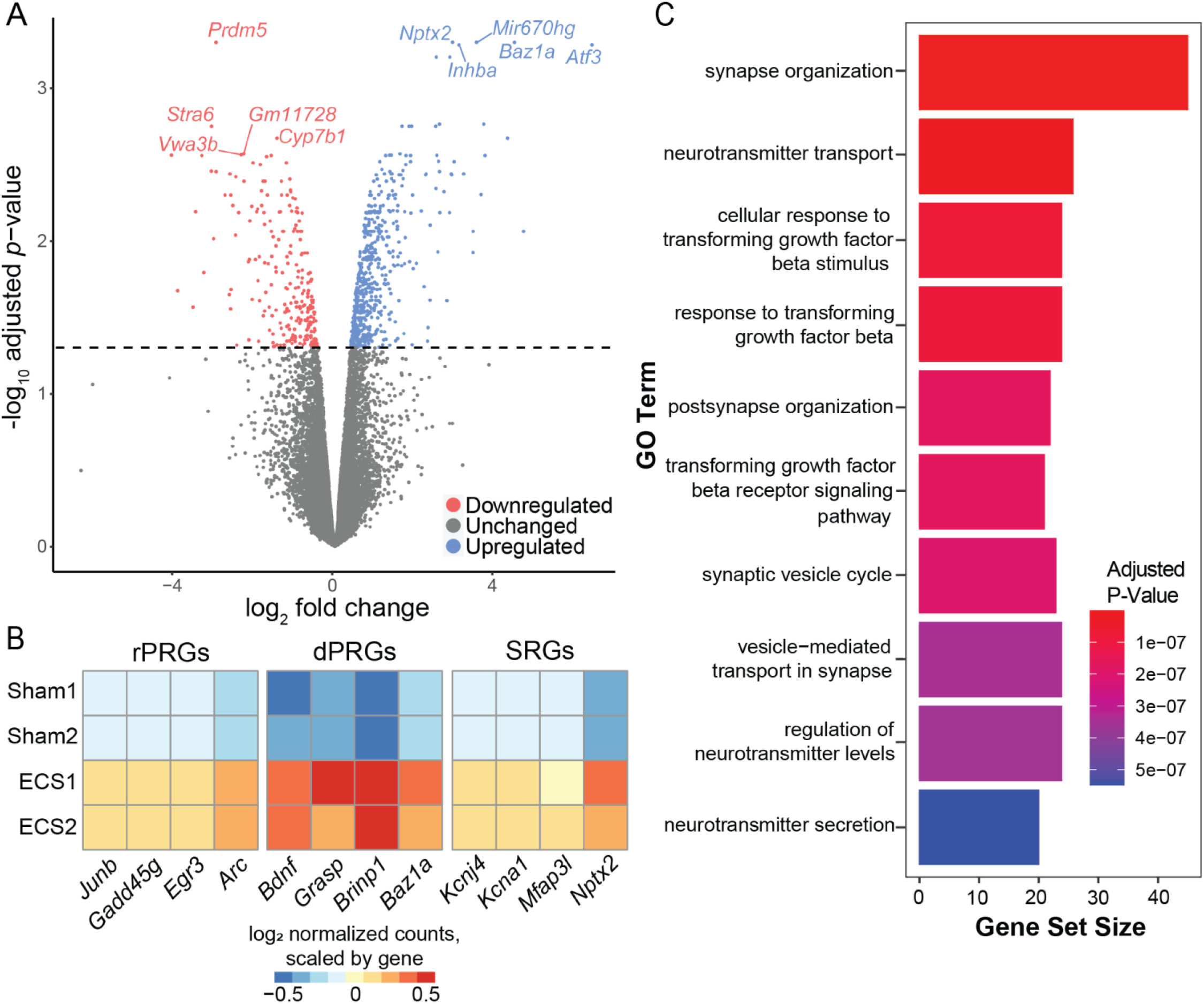
Differential expression results identifying activity-regulated genes (ARGs) associated with Sham versus ECS conditions. **(A)** Volcano plot illustrating DE analysis results with log_2_ fold change on the x-axis and FDR adjusted, -log_10_ transformed *p*-values on the y-axis. Genes colored blue are upregulated following ECS (FDR < 0.05, log_2_ fold change > 0), while genes colored red are downregulated (FDR < 0.05, log_2_ fold change < 0). Top 5 upregulated and downregulated genes are labeled. **(B)** Heatmap showing expression of 12 example activity-regulated genes across Sham (top 2 rows) and ECS replicates (bottom 2 rows), grouped by gene class: rapid primary response (rPRGs), delayed primary response (dPRGs), or secondary response (SRGs). **(C)** Bar plot visualizing GO term over-representation analysis results with bars corresponding to gene set size and bars colored by the FDR adjusted *p*-value.

### 3.3 Cell type-specific regulation of activity-regulated gene expression across distinct hippocampal neuron types

We also investigated cell type-specific differential expression between conditions (Sham and ECS) using pseudobulk DE analysis within each annotated cell type (**Figure S4; Table S4**). We combined GC.1 and GC.2 cell clusters for this analysis due to almost complete separation between GC nuclei by condition. These GC nuclei had the highest level of DE between conditions, compared with other groups (9,439 DE genes at 0.05 FDR threshold; 4,967 downregulated, 4,593 upregulated) (**Figure 3A-B**). Among other nuclei, CA3.1 (284 DE genes; 34 downregulated, 250 upregulated), PS.1 (179 DE genes; 13 downregulated, 166 upregulated), and CA1 (69 DE genes; 10 downregulated, 59 upregulated) had the next highest levels of differential expression (**Figure 3A-F**). Thus, we selected GC, CA3.1, CA1, and PS.1 clusters for further inquiry, focusing on particular genes with a positive log fold change, which we deemed upregulated genes (**Figure 3B-F**). There were 23 genes upregulated following ECS in all four clusters, including *Mir670hg*, a non-coding microRNA host gene and protein phosphatase *Ppm1h* (**Figure 3G-H**). We found 4,311 genes upregulated in GC and not CA1, CA3.1, or PS.1, including *Baz1a, Tll1*, and *Inhba* (**Figure 3I-K**). Nearly all upregulated DE genes following ECS for CA3.1, CA1, and PS.1 clusters were also upregulated in GC. However, a few genes were found to be exclusively upregulated in CA3.1 (32 genes), PS.1 (8 genes), and CA1 (2 genes). Additionally, 6 genes were upregulated in CA3.1, PS.1, and CA1, but not GC, including *Kdm2b* (**Figure 3L**). These results suggest that while much of the differential expression between conditions is driven by putative GC cells, there is still heterogeneity of DE genes across cell types in the HPC.

**Figure 3:**
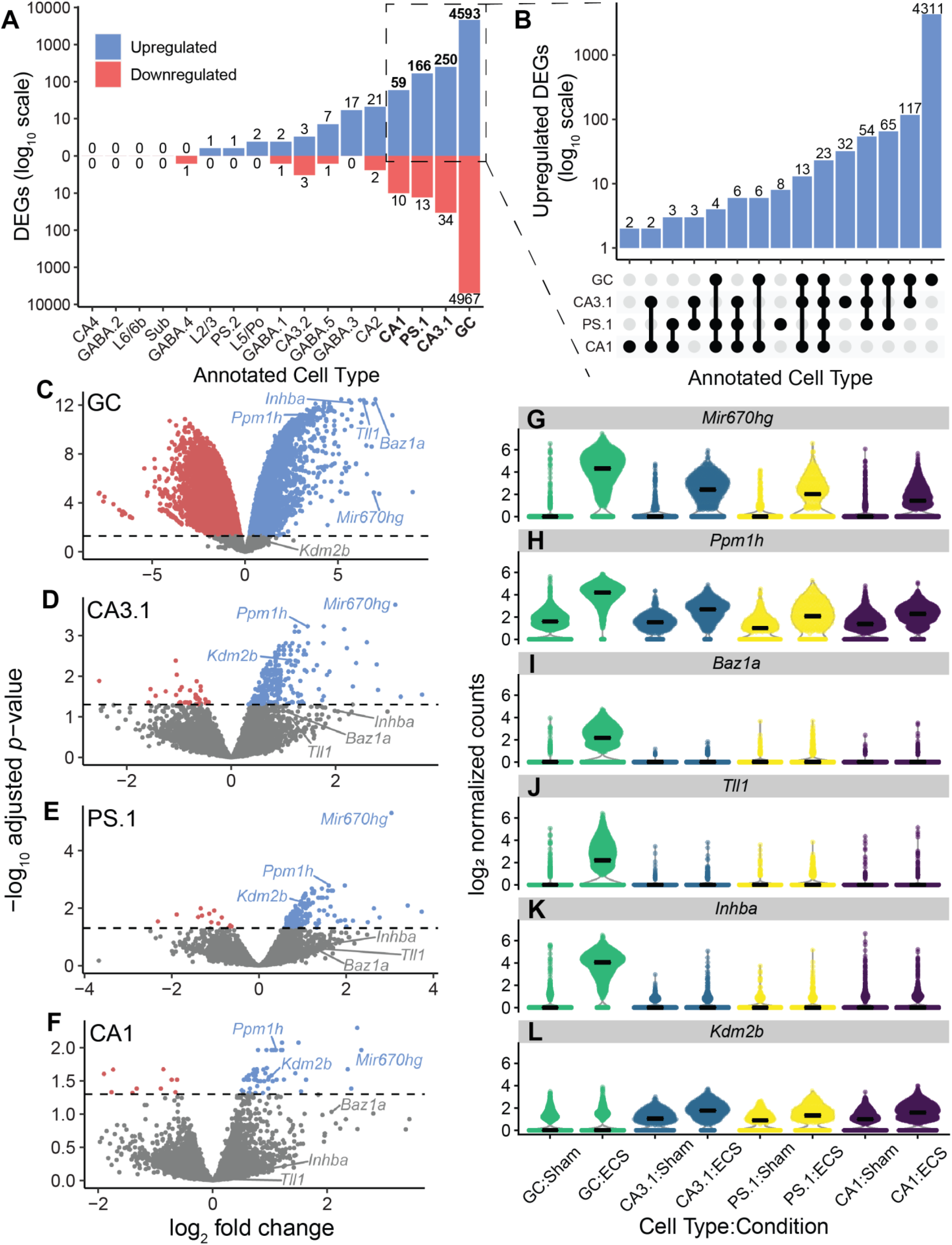
Differential expression results identifying cell type-specific activity-regulated genes associated with Sham versus ECS conditions. **(A)** Bar plot illustrating number of cell type-specific DE genes across conditions. Genes below zero (red) on the y-axis are downregulated (log_2_ fold change < 0) and genes above zero (blue) are upregulated (log_2_ fold change > 0). **(B)** UpSet plot (Conway et al., 2017; Ahlmann-Eltze, 2020) comparing upregulated DE genes across GC, CA3.1, PS.1, and CA1 annotations. Annotated cell types are shown on the left of the modified x-axis, and x-axis elements represent all possible intersections between DE genes between conditions for these cell types. The y-axis is shown on a log_10_ scale and indicates the number of genes at each intersection (e.g. there are 23 genes that are DE between conditions for all cell types). **(C)** Volcano plot visualizing DE analysis results for GC nuclei. Points colored blue are DE genes with log_2_ fold change > 0 and points colored red are DE genes with log_2_ fold change< 0. Labeled genes are illustrated in panels F-H. **(D)**Same as C, but for CA3.1 nuclei. **(E)** Same as C and D, but for PS.1 nuclei. **(F)** Same as C-E, but for CA1 nuclei. **(G)** Violin plot showing differential expression of *Mir670hg* in GC, CA3.1, PS.1, and CA1 nuclei across both conditions. **(H)** Same as G, but for *Ppm1h*. **(I)** Same as G&H, but for *Baz1a*. **(J)** Same as G-I, but for *Tll1*. **(K)** Same as G-J, but for *Inhba*. **(L)** Same as G-K, but for *Kdm2b*.

### 3.4 Functional enrichment analysis of cell-type specific activity-regulated gene expression

To assess biological processes that may be altered in specific cell types in response to ECS, we used gene set enrichment analysis (GSEA) to probe our cell type-specific DE analysis results. For each cell type, using a *t*-statistic calculated from DE analysis results for all genes, we ran GSEA across all GO term gene sets for Biological Process (BP), Cellular Component (CC), and Molecular Function (MF) domains. Across all cell types, we found enrichment of 1,029 GO terms (814 BP, 118 CC, 97 MF). These terms were enriched in specific cell types, with GC having the most enriched terms (650 BP, 80 CC, 68 MF), followed by CA3.1 (359 BP, 70 CC, 50 MF), PS.1 (414 BP, 64 CC, 48 MF), and CA1 (363 BP, 61 CC, 45 MF) (**Figure S5A**).

Due to the large size and number of gene sets associated with GO terms, GSEA results often contain redundancies that render interpretation difficult. To address this issue, we performed semantic similarity analysis to cluster GO terms by similarity. We pooled all terms enriched in at least one cell type by GO domain. This analysis identified 28 BP clusters, 18 CC clusters, and 20 MF clusters (**Figure S5B-D; Figure S6; Table S5**). For example, one CC cluster was entirely composed of terms related to synapses, such as “presynapse,” “postsynaptic density,” and “glutamatergic synapse” (**Figure 4A**). Upregulated genes associated with terms in this cluster had various roles, including receptor subunits (*Grin2a*), molecular scaffolds (*Dlg4, Grasp*), and synaptic vesicle proteins (*Sv2b, Svop*). Some, such as *Arc* and *Nptx2*, are known ARGs. These genes had heterogeneous levels of DE across annotated cell types (**Figure 4B**). Another cluster was enriched for groups of terms describing signaling pathways in response to different stimuli, such as “response to organic cyclic compound,” “response to cytokine,” and “response to growth factor” (**Figure 4C**). Genes associated with these clusters included genes involved with modulation of growth factor signaling pathways (*Ntrk2, Bdnf*), but also regulators of inflammation and immune signaling (*Ptgs2, Tgfb2, Smad3*), which were also differentially upregulated across cell types (**Figure 4D**). Among MF clusters, one contained only terms associated with ubiquitin or ubiquitin-like ligase or transferase activity (**Figure 4E**). Indeed, gene sets for these terms mostly included different E2 ubiquitin enzymes (*Ube2g1, Ube2q2*), E3 ubiquitin ligases (*Cop1, Nedd4l, Mib1*), E3 ubiquitin ligase or genes that recruit ubiquitination enzymes (*Bcor*). These again displayed heterogeneous patterns of expression (**Figure 4F**).Finally, one cluster was enriched with groups of MF terms associated with kinase activity (**Figure 4G**), with genes coding for different kinases (such as *Plk2, Akap13, Mapk4, Hunk*, and *Itpk1*). Together, these results demonstrate that ECS alters expression of genes associated with various biological processes across multiple cell types, with a particularly strong impact on dentate GCs.

**Figure 4:**
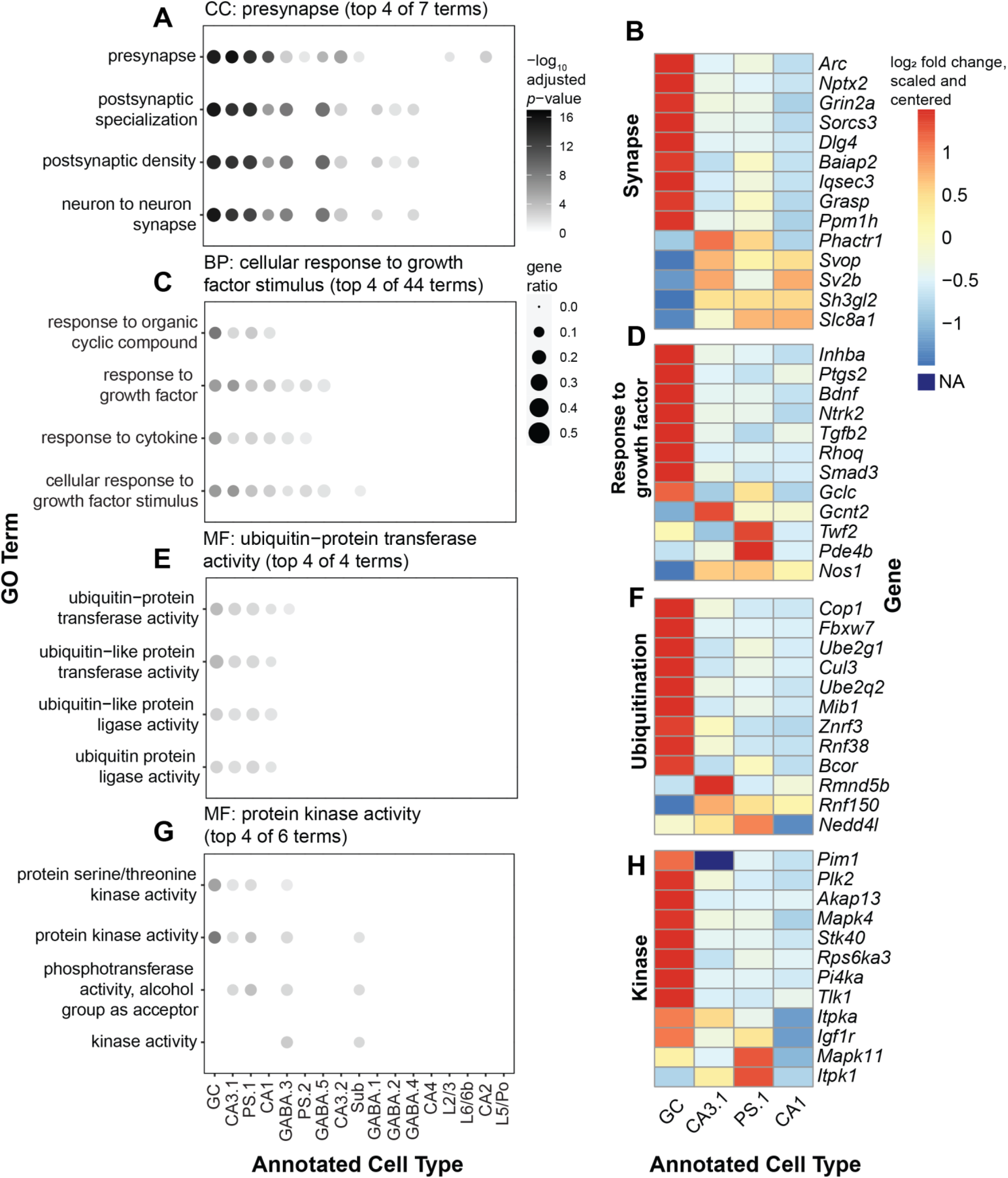
Cell type-specific Gene Set Enrichment Analysis (GSEA) in Sham versus ECS conditions. **(A)** Plot showing enrichment of clustered Cellular Component (CC) GO terms obtained by semantic similarity analysis of cell type-specific GSEA results. Example cluster grouped 10 terms associated with synapses; top 6 are shown. Circles are colored by -log10 FDR-adjusted p-value and sized by the gene ratio for each term (Methods). **(B)** Heatmap showing cell type-specific differential expression of genes associated with terms in the corresponding CC GO term cluster. Heatmap is colored by -log_2_ fold change, scaled by gene and centered at 0. **(C)** Same as A, but for Biological Process (BP) GO terms. Example cluster grouped 40 terms associated with cell signaling pathways; top 6 are shown. **(D)** Heatmap showing cell type-specific differential expression of genes associated with terms in the corresponding BP GO term cluster. **(E)** Same as A and C, but for Molecular Function (MF) GO term cluster grouping 4 terms associated with ubiquitination activity. **(F)** Heatmap showing cell type-specific differential expression of genes associated with terms in the corresponding MF GO term cluster. **(G)** Same as A, C, and E, but for MF GO term cluster grouping 6 terms associated with kinase activity; top 4 are shown. **(H)** Heatmap showing cell type-specific differential expression of genes associated with terms in the corresponding MF GO term cluster.

### 3.5 Non-negative matrix factorization identified both cell type-specific and ECS-induced gene expression profiles

Dimensionality reduction techniques can be powerful analysis methods for novel discovery in sequencing data. For example, the `CoGAPS` Bioconductor package uses Bayesian non-negative matrix factorization (NMF) to identify gene expression patterns, including an application for snRNA-seq or scRNA-seq data. Briefly, `CoGAPS` decomposes a gene count matrix into two matrices corresponding with sources of variation across genes and cells, respectively. These matrices can then be used to define gene expression patterns within the original dataset, which might correlate with technical artifacts, cell types, or biological processes (Stein-O’Brien et al.,2019). We asked if this approach would identify potential activity-regulated gene expression signatures associated with ECS in our dataset. `CoGAPS` identified 57 distinct gene expression patterns (**Figure 5A**). Most of these patterns show good correspondence with the cell type clusters as annotated in Sections 3.1 and 3.2. For instance, Patterns 52 and 57 were both enriched in CA3.1 neurons. Other patterns had a lower level of correlation across multiple clusters, such as Pattern 26, which is associated with several RHP clusters. Using the `patternMarkers()` function provided in the `CoGAPS` package, we identified sets of marker genes for each of these patterns (**Table S6**).

**Figure 5:**
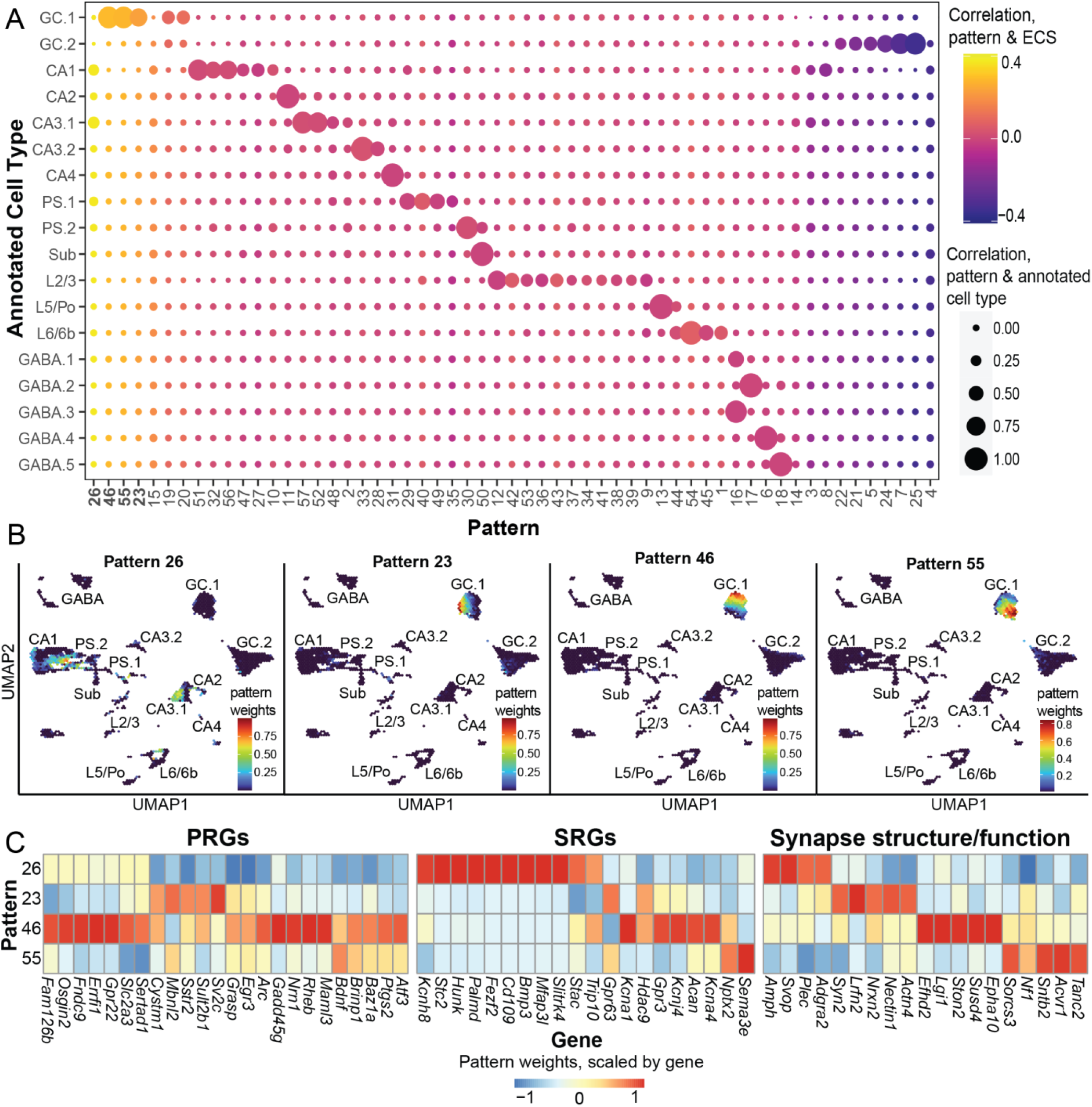
Identification of gene expression patterns using scCoGAPS. **(A)** Balloon plot comparing CoGAPS patterns, annotated cell types, and ECS condition. Circles are colored by correlation between CoGAPS patterns and ECS and sized by correlation between CoGAPS patterns and annotated cell type. **(B)** UMAP representations of the data illustrating the 4 patterns most highly correlated with ECS. Plots are colored by pattern weights for patterns 26, 23, 46, and 55, respectively. **(C)** Heatmap illustrates differences in gene expression across patterns 26, 23, 46, and 55. Heatmap is colored by pattern weights for each gene, scaled by gene and centered at 0. Genes are classified as primary response genes, secondary response genes, or genes involved with synaptic structure and function.

Four patterns (26, 23, 46, and 55) were more highly correlated with ECS than others. Of these, pattern 26 was mostly correlated with pyramidal cell clusters, namely CA1, CA3.1, and PS.1. Three patterns (23, 46, and 55) were enriched in GC.1 cells in a largely non-overlapping fashion (**Figure 5B**). Using marker genes for these patterns, we found heterogeneity of expression of known ARGs, including PRGs and SRGs, along with genes involved with synapse structure and function (**Figure 5C**). Taken together, these results suggest the presence of multiple continuous, activity-regulated gene expression patterns in mouse hippocampal neurons.

## 4 Discussion

Neural plasticity in the HPC is critical for mediating many biological processes that underlie learning and memory. These processes include structural and functional changes in HPC cells that regulate various forms of plasticity. This plasticity depends on waves of activity-regulated transcription. The early waves consist of primary response genes and immediate early genes, which are used to mark activated populations of neurons. These waves are followed by transcription of secondary response genes. There is also evidence that these ARG waves functionally contribute to synaptic plasticity underlying learning and memory. However, many questions remain about ARG transcription across cell types. For example, it is not known whether activity-induced increases are due primarily to increased expression of activity-regulated transcripts in a small population of recruited cells, or alternatively whether cells across multiple sub-populations are broadly recruited. Our data show that the transcriptional representation of activity is robust, but differs across cell types.

Here, we used snRNA-seq to identify activity-regulated genes and gene expression patterns across major neuronal cell types in the mouse HPC in response to ECS. We identified large ECS-induced changes in gene expression in dentate GCs and moderate, yet heterogeneous, differential expression across populations of glutamatergic neurons outside the dentate. Our analyses revealed that > ⅓ of detected genes were differentially expressed between conditions in dentate GCs. Dentate GCs are under strong tonic inhibition from GABAergic neurons (Jung and McNaughton, 1993; Li et al., 2013; Engin et al., 2015; Pardi et al., 2015) and have an elevated threshold for excitation (Krueppel et al., 2011), which contributes to their sparse activation under normal conditions and renders them uniquely positioned to gate strong excitatory input (Hainmueller and Bartos, 2020;Leutgeb et al., 2007). However, brain-wide stimulation, such as ECS, can override this inhibition, subsequently altering downstream network activity in the HPC. Our work is consistent with previous rodent microarray studies of HPC as well as laser microdissected dentate gyrus, which found substantial ECS-induced changes in genes associated with neurotransmission, neurotrophic signaling, angiogenic signaling, and other growth factor signaling pathways implicated in neuroprotection. Induction of these molecular pathways may contribute to the ability of ECS to stimulate dendritic growth and arborization in immature granule cells and enhanced excitability in mature GCs (Imoto et al., 2017; Ueno et al., 2019; Marín-Burgin et al., 2012). Together, the data support the notion that ECS-induced activity may contribute to a state shift in which normally quiescent GCs are molecularly primed for growth, development, and survival.

ECS-driven gene expression changes were also found in other excitatory populations, with cells from the CA3.1 cluster showing the highest level of differential expression, followed by CA1, and then CA2 cells. However, ECS had little to no effect on gene expression in CA3.2 and CA4 clusters. There are several reasons why excitatory neuron populations may be differentially impacted by ECS. First, the neuroanatomical organization of the molecular circuit may drive these changes. Specifically, GCs synapse primarily onto CA3 neurons, which project to CA1 and CA2 neurons via Schaffer collateral axons. Since CA3.1 and CA1 cells are synaptically downstream of GCs, differential expression being strongest in GCs may simply reflect the sequence of current flow. Second, HPC neuron subtypes exhibit different levels of excitability and synaptic plasticity that are likely reflected in their gene expression profiles. For example, CA2 neurons are less susceptible to long term potentiation (LTP) compared with CA1 and CA3 neurons (Carstens and Dudek, 2019). Third, gene expression, connectivity, and electrophysiological characteristics of neurons within the same HPC subfield can vary based on anatomical position (Cembrowski and Spruston, 2019). Indeed, ventral CA1 neurons tend to be more excitable than dorsal CA1 neurons (Dougherty et al., 2012). CA3 neurons also have very different connection patterns, firing properties, and roles in memory formation along the proximodorsal and superficial-deep axes (Lee et al., 2015; Sun et al., 2017; Hunt et al., 2018). In our dataset, CA3.2 neurons may correspond with deep CA3 neurons as reflected by enriched expression of *St18*, a marker for a rare subtype of very deep CA3 pyramidal cells, whereas CA3.1 cells are likely more superficial due to much lower expression of this marker gene (Bienkowski et al., 2018; Thompson et al., 2008).Thus, among HPC pyramidal neurons, cell type-specific transcriptional responses to ECS may be explained by the convergence of molecular, cellular, and circuit-level patterns governing anatomical position, neuronal excitability, and network connectivity.

Our study also revealed novel biology surrounding the subiculum and prosubiculum, which have been understudied due to challenges in anatomical and molecular classification. We observed heterogeneity in transcriptional response to ECS in clusters that map to subiculum and prosubiculum; specifically, PS.1 neurons showed moderate levels of differential expression between conditions, while PS.2 and Sub clusters showed little to none. These disparities may stem from anatomical locations of these clusters. PS.1 neurons may be more superficial (*Nos1+*) and ventral (*Dcn*+), while PS.2 may be deeper (*Nos1-, Cntn6*+) and more dorsal (*Ndst4*+) (Cembrowski et al., 2016a; Habib et al., 2016; Ding et al., 2020). Ding et al. classify the ventral subiculum as the ‘ventral prosubiculum’ except for the most caudal portion of the subicular complex, based on the expression of PS-specific genes (*Ntng2*) and lack of expression of subiculum-specific genes (*Nts*, *Fn1*) in all but this most caudal portion (Ding et al., 2020). Similarly, we observe PS-specific genes in both PS.1 and PS.2, notably *Ntng2*, and subiculum-specific genes only in our Sub cluster, namely *Nts* and *Fn1*. A recent bulk RNA-seq study stimulated the perforant path to acutely induce seizures, and found higher levels of differential expression in ventral versus dorsal subiculum (Aoyama et al., 2022). Our finding that PS.1 cluster is regulated by ECS lends support to this result, since PS.1 likely corresponds with what Aoyama et al. (2022) described as ventral subiculum because PS.1 is enriched for ventral-specific marker genes. Prosubiculum projection neurons, particularly those in the ventral portion, target a broader range of subcortical structures than subiculum projections, including the amygdala, ventral striatum, and hypothalamus (Ding et al., 2020). Subiculum neurons also have different firing properties based on their projection targets and anatomical location (Cembrowski and Spruston, 2019; Kim and Spruston, 2012; Graves et al., 2012; Böhm et al., 2015). Taken together, heterogeneity of ECS-induced gene expression changes across subicular and prosubicular clusters may reflect differential recruitment across the dorsal-ventral axis that impacts HPC output to critical target regions.

A unique technical aspect of our study is that we utilized two distinct matrix factorization approaches, PCA and NMF, that decompose high-dimensional data into a lower-dimensional set of factors. In contrast to PCA, which aims to identify ordered factors that explain the most amount of variation, NMF aims to identify factors that represent independent, additive features in the data. NMF is not utilized as often as PCA in bioinformatics because it tends to be much slower and more computationally expensive. However, a recently published NMF implementation greatly increases its speed, which may enable more widespread usage (DeBruine et al.,2021). In our study, most NMF factors corresponded well with previously annotated cell types. However, we also found the presence of several factors associated with ECS, including three associated with granule cells and one associated with various pyramidal cell types, namely CA3.1, and PS.1, and CA1 neurons. These four factors may represent distinct, cell type-specific, activity-regulated patterns of gene expression. Earlier studies indicate that NMF factors identified within gene expression data may correspond with independent sources of variation, which may be biological (e.g. cell type, cell state, biological process) or technical (e.g. batch effects). These patterns can be thought of as continuous gene expression signatures that represent complex biological processes (Stein-O’Brien et al., 2018). This may be a more nuanced and informative approach to assessing the impact of experimental conditions than the presence or absence of sets of marker genes. Previous studies have also taken an NMF-based approach to identify patterns of activity-induced gene expression in scRNA-seq data, particularly in cortical neurons (Kotliar et al., 2019). Additionally, these biological processes may be shared across datasets, particularly other sequencing datasets in the hippocampus. Future studies can thus leverage NMF factors identified in this study to identify activated cell populations within other transcriptomic datasets.

ECS is a translational laboratory model for electroconvulsive therapy (ECT) that can provide clinical insights by revealing the neurobiological mechanisms underlying antidepressant efficacy. While ECT remains one of the most rapid and effective treatments for major depression, it requires repeated administration of anesthesia, and can have adverse cognitive side effects (UK ECT Review Group, 2003; Kho et al., 2003). Several lines of evidence suggest that changes in gene expression downstream of neural activity induction after ECT influence circuit function that controls behavior (Bolwig, 2011). Understanding the molecular pathways that are initiated in discrete cell populations is important because this information can be used to develop more specific ECT-mimetics or to refine brain stimulation protocols. Defining the molecular profile of dentate GCs that are particularly responsive to ECS can facilitate future work to selectively target these cells. We also identified enrichment of molecular signaling pathways related to immune, growth factor, and cytokine signaling, which may be translationally relevant given emerging findings linking immune dysregulation and inflammation with risk for depression (Goldfarb et al., 2020; Miller and Raison, 2016; Dantzer et al., 2008; Beurel et al., 2020) While immune response to ECS may result in increased inflammation, we also found enrichment of TGF-beta and related growth factor signaling pathways. In neurons, TGF-beta serves important roles in neurodevelopment, axon guidance, synaptic plasticity, intracellular signaling, and cell survival (Meyers and Kessler, 2017; Mitra et al., 2022; Krieglstein et al., 2011), and has anti-inflammatory and neuroprotective properties. Some ECS-induced genes involved with immune signaling and inflammation may also have multifaceted roles in ECT. For instance, *Ptgs2* has a well-known role in inflammatory response and may be involved in processes important to the antidepressant effects of ECT, such as synaptic plasticity and modulation of synaptic transmission (López and Ballaz, 2020; Stark and Bazan, 2011; Cowley et al., 2008), and the adverse cognitive side effects of ECT (Andrade et al., 2008). More research is necessary to parse the nuanced roles of different genes and signaling pathways involved in ECT, and our findings can help to inform hypothesis generation in these studies.

Our study has a number of limitations. Specifically, the low nuclei number in some of the smaller clusters may limit our power to detect differentially expressed genes. In addition, our study was focused on profiling neuronal subtypes and their response to ECS and we therefore did not profile glia, which are functionally impacted by ECS (Wennström et al., 2003; Jansson et al., 2009; Steward, 1994; Ongür et al., 2007). Finally, our experimental paradigm only included a single ECS session, and did not investigate chronic gene expression changes. Since typical courses of ECT involve multiple sessions (Charlson et al., 2012), similar studies with repeated ECS administration will be important. In summary, these data contribute to a more complete understanding of ARG expression in HPC, which is crucial to translate ECS research from animal models to treatments for human disease. To facilitate access of the data to the community for further exploration, we provide our data as a resource in an accessible web-based format along with all code for reproducible analysis of the data.

## Acknowledgements

We thank members of the Martinowich, Maynard, and Hicks laboratories for discussions and comments on the manuscript. We thank the Johns Hopkins University Sidney Kimmel Comprehensive Cancer Center (SKCCC) Flow Cytometry Core and the Johns Hopkins University Single Cell and Transcriptomics Core for supporting snRNA-seq experiments. This project was supported by the Lieber Institute for Brain Development, National Institutes of Health awards R21MH118725 (KM), R01MH105592 (KM), and Chan Zuckerberg Initiative DAF, an advised fund of Silicon Valley Community Foundation CZF2019-002443 (SCH). KRN was supported by the National Institutes of Health award R25GM109441. All funding bodies had no role in the design of the study and collection, analysis, and interpretation of data and in writing the manuscript.

## Data Availability Statement

Raw FASTQ sequence data files will be made available from SRA. The code to reproduce the data analyses and figures, along with 2 `SingleCellExperiment` objects containing the 15,990 HPC nuclei and the 17,804 nuclei before thalamic neuron removal, are available on GitHub (https://github.com/Erik-D-Nelson/ARG_HPC_snRNAseq). Data is also available in a web-based iSEE app format at https://libd.shinyapps.io/2022_HPC_ARG/ (Rue-Albrecht et al., 2018).

## Author Contributions

Conceptualization: KRM, KM

Data Curation: MNT, EDN

Formal Analysis: EDN

Funding acquisition: KM, KRM, SCH

Investigation: KRM, KRN, MNT

Project Administration: KM, SCH

Software: HRD, LCT

Supervision: KRM, KM, SCH

Writing – original draft: EDN, KRM, KM, SCH

Writing – review & editing: EDN, KRM, KM, SCH

## Supplementary Information

**Table S1: Marker gene statistics for all clusters.** Data is provided as an .xlsx file. Each sheet within this Excel workbook contains marker gene statistics for each cluster, obtained by performing pairwise *t*-tests between all pairs of clusters for every gene. Columns include p-values, FDR adjusted p-values, summary logFC (combined logFC across all other clusters), and log FC values.

**Table S2:**

**Statistics from initial (non-cell type specific) differential expression (DE) analysis between the Sham and ECS conditions.** Data is provided as an .xlsx file. Columns include Ensembl ID for each gene (“ID”), gene symbol, gene type (“gene_biotype”, one of “protein_coding”, “lincRNA”, or “antisense”), log_2_ fold change (“logFC”), log_2_ counts per million reads (“logCPM”), *F*-statistic, *p*-values, and FDR-adjusted *p*-values.

**Table S3:**

**Over-representation analysis (ORA) results for enrichment of biological process (BP) GO gene sets within identified DE genes between conditions from sample-level DE analysis.** Data is provided as an .xlsx file. Columns include GO ID (“ID”), GO term description (“Description”), gene ratio for each term (“GeneRatio”), ratio of all detected genes associated with each term to all detected genes (“BgRatio”), *p*-values, FDR adjusted *p*-values (“p.adjust”), *q*-values, Ensembl gene IDs, and set size of DE genes associated with a given term (“Count”).

**Table S4:**

**Statistics from cell type specific differential expression (DE) analysis between the Sham and ECS conditions.** Data is provided as an .xlsx file. Each sheet within this Excel workbook contains DE analysis statistics for each cell type. Columns for each sheet are the same as Table S2.

**Table S5:**

**Semantic similarity clustering results for GSEA results.** Data is provided as an .xlsx file. Each sheet contains clustering results for all enriched GO terms within each domain (BP, CC, MF). Columns include GO ID (“go”), cluster number, parent GO ID (“parent”), cluster similarity score (“parentSimScore”), mean -log_10_ adjusted *p*-value across all cell types (“score”), gene set size, and GO term description (“term”).

**Table S6: Lists of marker genes for CoGAPS patterns.** Data is provided as an .xlsx file. Each sheet contains a list of marker genes for a different CoGAPS pattern.

**Figure S1:**
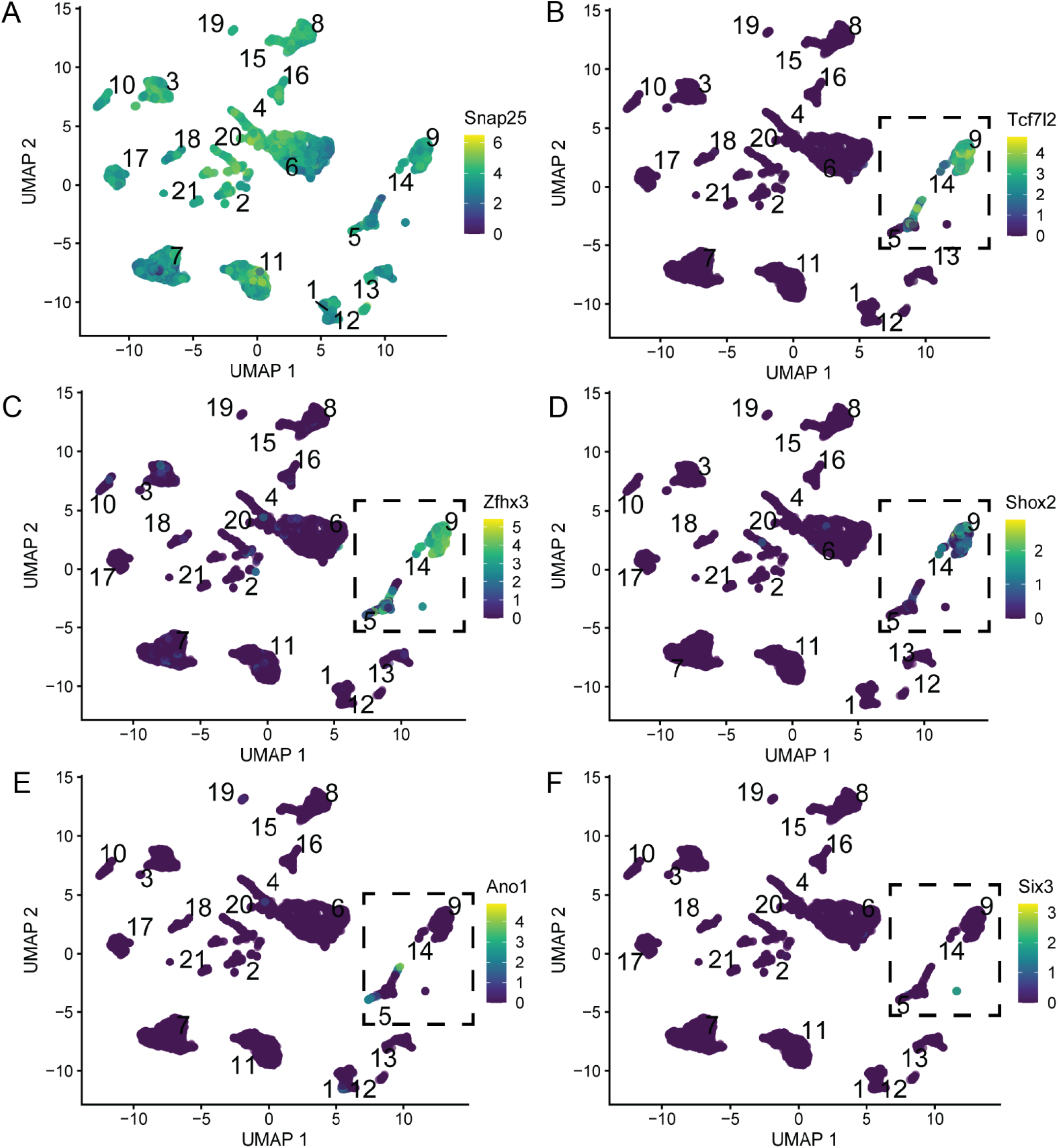
Identification of thalamic and epithalamic nuclei. **(A)** UMAP representation of dataset prior to thalamic cluster removal. The plot is labeled by preliminary clustering and colored by *Snap25* expression (in log_2_-normalized counts). All clusters express *Snap25,* a marker gene for neurons. **(B)** UMAP colored by expression of *Tcf7l2*, a thalamus and epithalamus-specific transcription factor. Expression is limited to clusters 5, 9, and 14. **(C)** UMAP colored by expression of *Zfhx3*, a thalamus-specific transcription factor. **(D)** UMAP colored by expression of *Shox2*, a thalamus-specific marker gene. **(E)** UMAP colored by expression of *Ano1*, an epithalamus-specific marker gene. **(F)** UMAP colored by expression of *Six3*, a thalamus-specific marker gene.

**Figure S2:**
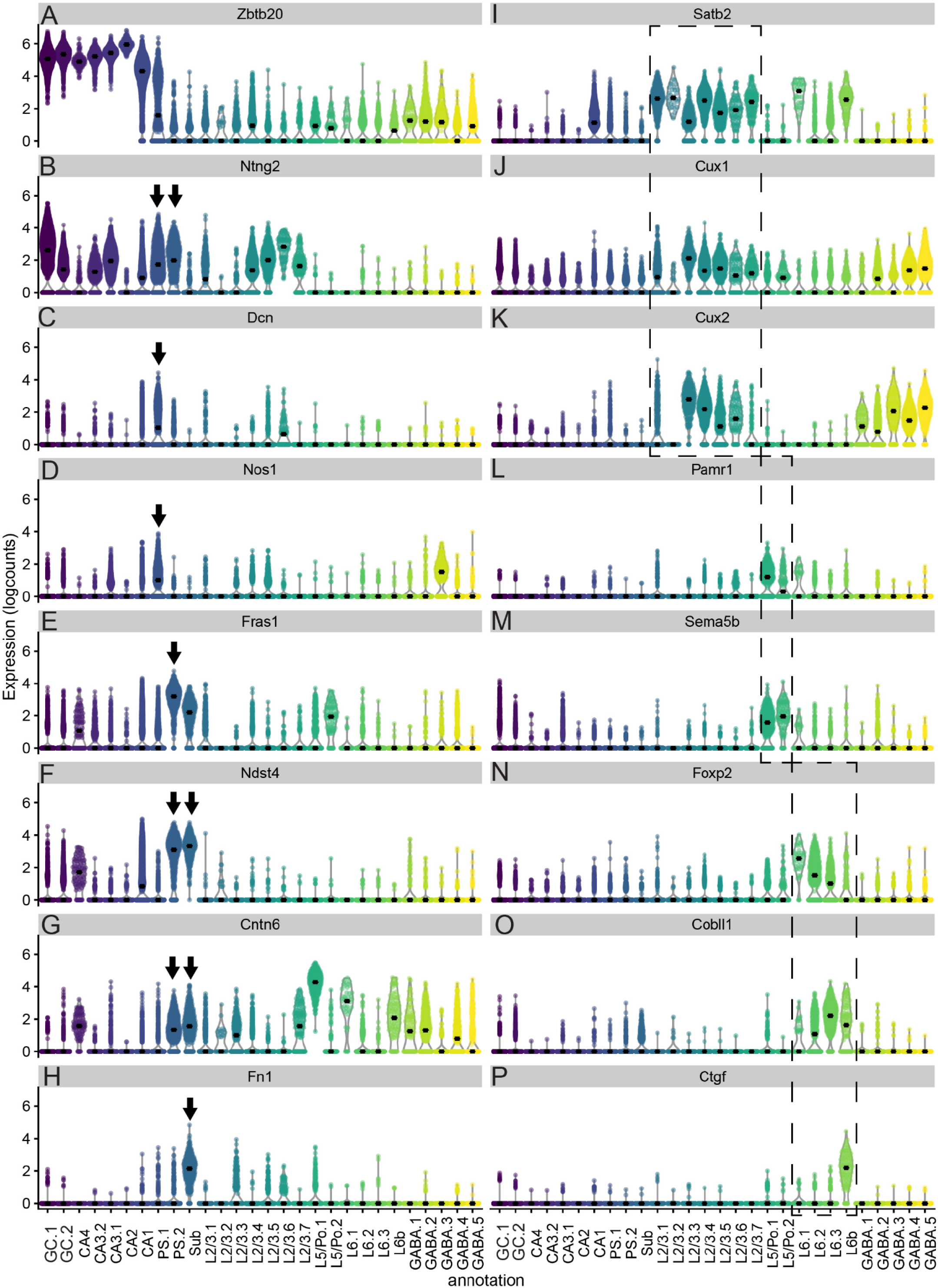
Additional characterization of clusters in mouse hippocampus. Violin plots showing expression (in logcounts) of various genes across 28 annotated clusters identified by graph-based clustering. **(A)** All clusters from dentate gyrus and cornu ammonia (CA) regions express high levels of *Zbtb20*. **(B)** Both prosubiculum (PS) clusters (arrows) express *Ntng2*, known to be present in PS but not subiculum (Sub). **(C-D)** PS.1 expresses *Dcn* and *Nos1*, known markers of ventral and superficial PS, respectively. **(F)** *Fras1* is a marker gene for PS.2 (arrow). **(F)** PS.2 and Sub clusters (arrow) express *Ndst4*, a known marker gene for dorsal Sub/PS. **(G)** PS.2 and Sub (arrows), but not PS.1, express *Cntn6*, which is known to be present in deep PS and Sub regions, but not superficial PS. **(H)** The Sub cluster (arrow), but not PS.1 and PS.2, expresses Fn1, a well-known Sub marker. **(I-K)** Expression of three known cortical/retrohippocampal region (RHP) layer 2/3 marker genes, *Satb2, Cux1*, and *Cux2*. Note enrichment in 7 L2/3 clusters (dotted rectangle), which are grouped together as L2/3 for downstream analyses. **(L-M)** Expression of *Pamr1* and *Sema5b*, which are expressed in RHP layer 5/polymorphic (Po) layer and cortical layer 5. Note enrichment in 2 L5/Po clusters (dotted rectangle), which are grouped together as L5/Po for downstream analyses. **(N-P)** Expression of three known cortical/RHP layer 6/6b marker genes, *Foxp2, Cobll1*, and *Ctgf*. Note enrichment in 3 L6 and 1 L6b clusters (dotted rectangle), which are grouped together as L6/6b for downstream analyses.

**Figure S3:**
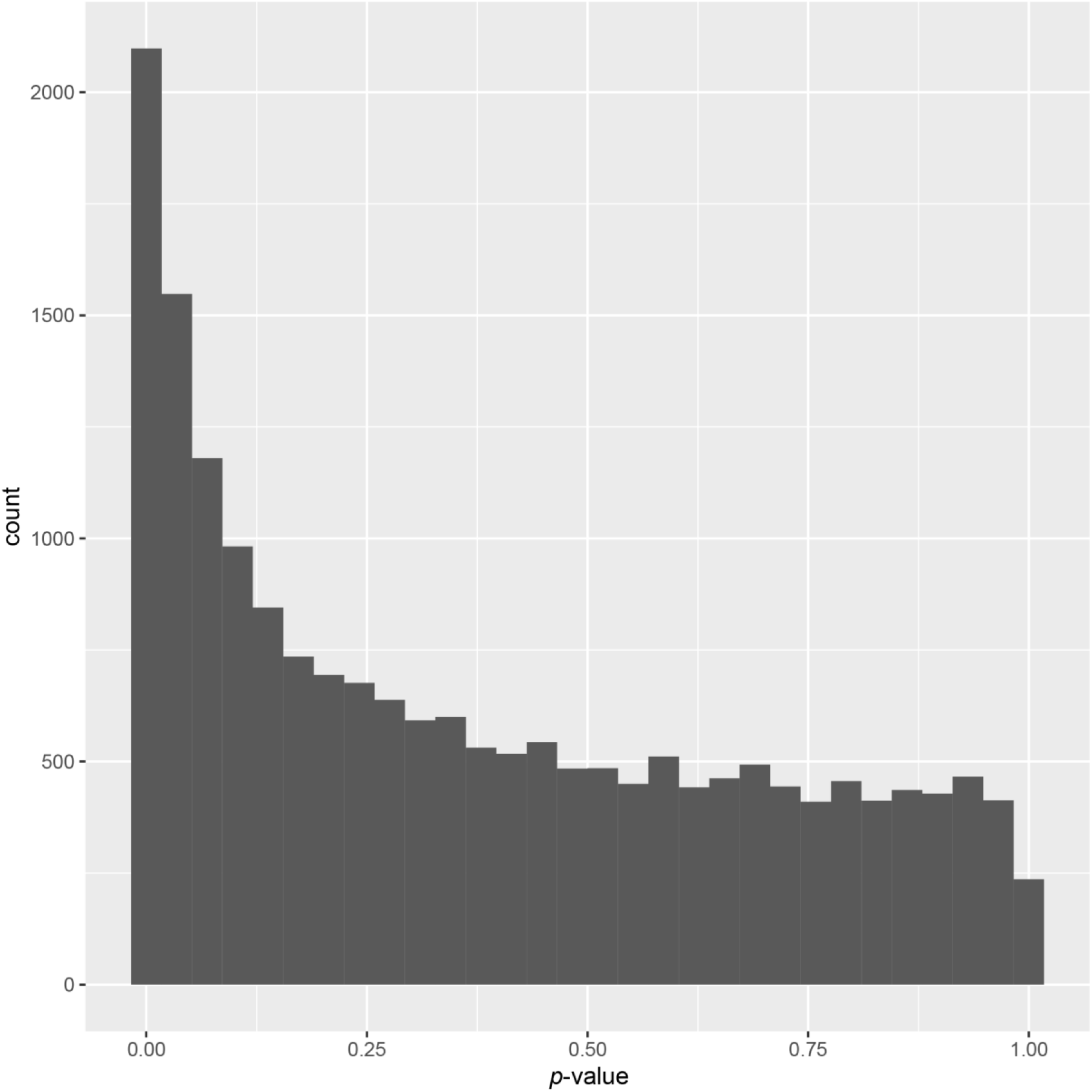
Histogram of *p*-values for all genes from sample-level DE analysis. DE analysis performed with edgeR after pseudobulking the data across all nuclei within each sample (N=2 Sham, N=2 ECS). Note the single peak of *p*-values centered around 0. The rest of the distribution is relatively uniform, but slopes down from left to right (from 0 to 1). This indicates both an abundance of alternative hypotheses for this analysis and that assumptions made in our modeling approach and statistical tests during DE analysis are valid for our dataset’s distribution (Breheny et al., 2018).

**Figure S4:**
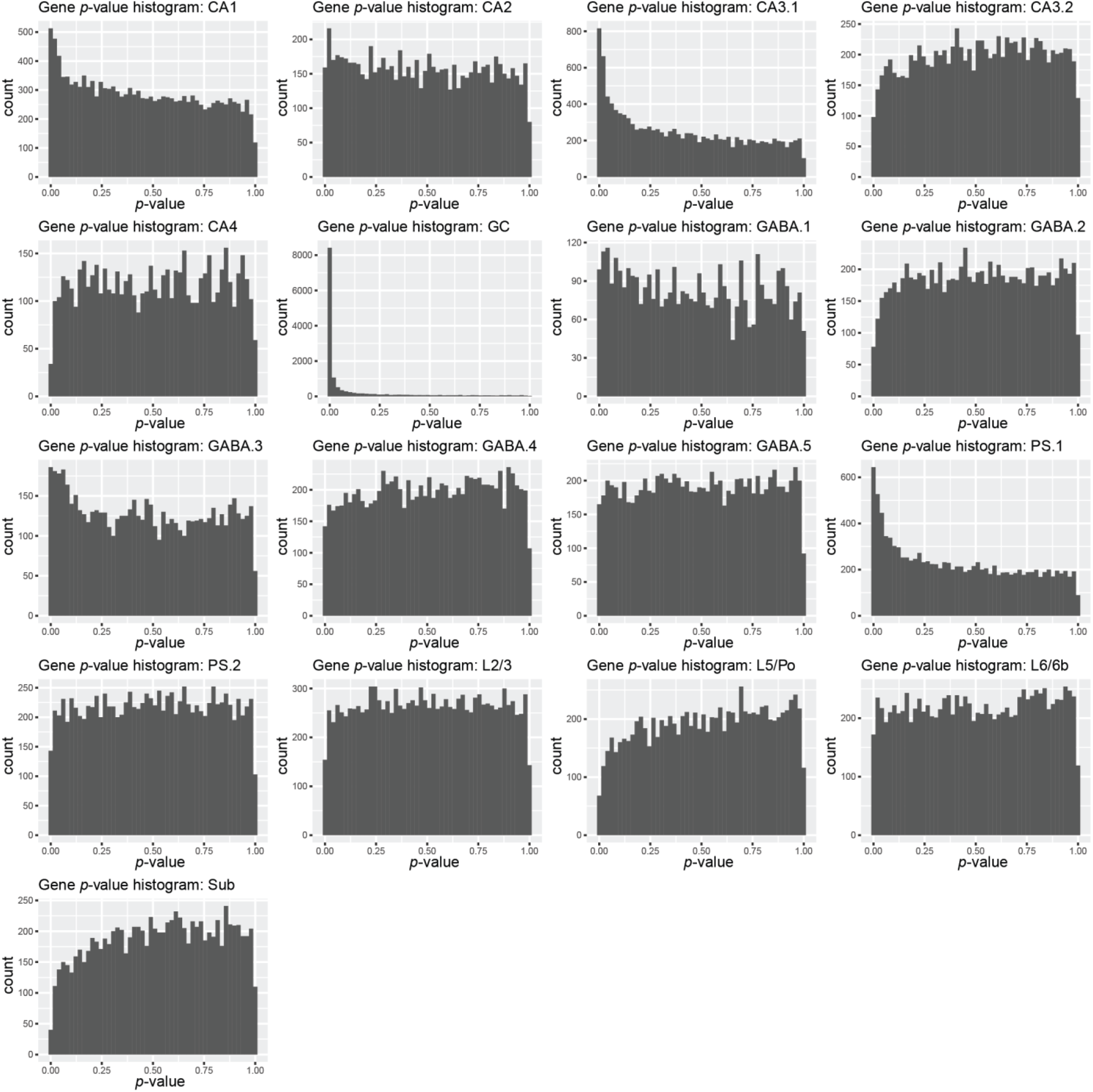
Histograms of *p*-values for each cell type from cell type-specific DE analysis. Same as Figure S3, but for cell-type specific DE analyses. Note four cases (CA1, CA3.1, GC, and PS.1) with peaks of *p*-values centered around 0, similar to the distribution in Figure S3. The distribution of *p*-values for GC has the largest peak, followed by CA3.1, PS.1, and CA1. All other cases are relatively uniform. This indicates a large number of non-null hypotheses for GC, CA3.1, PC.1, and CA1 cases, with other cell types having mostly null hypotheses. Once again, these plots indicate that assumptions made in our modeling approach and statistical tests during DE analysis are valid for our dataset’s distribution.

**Figure S5:**
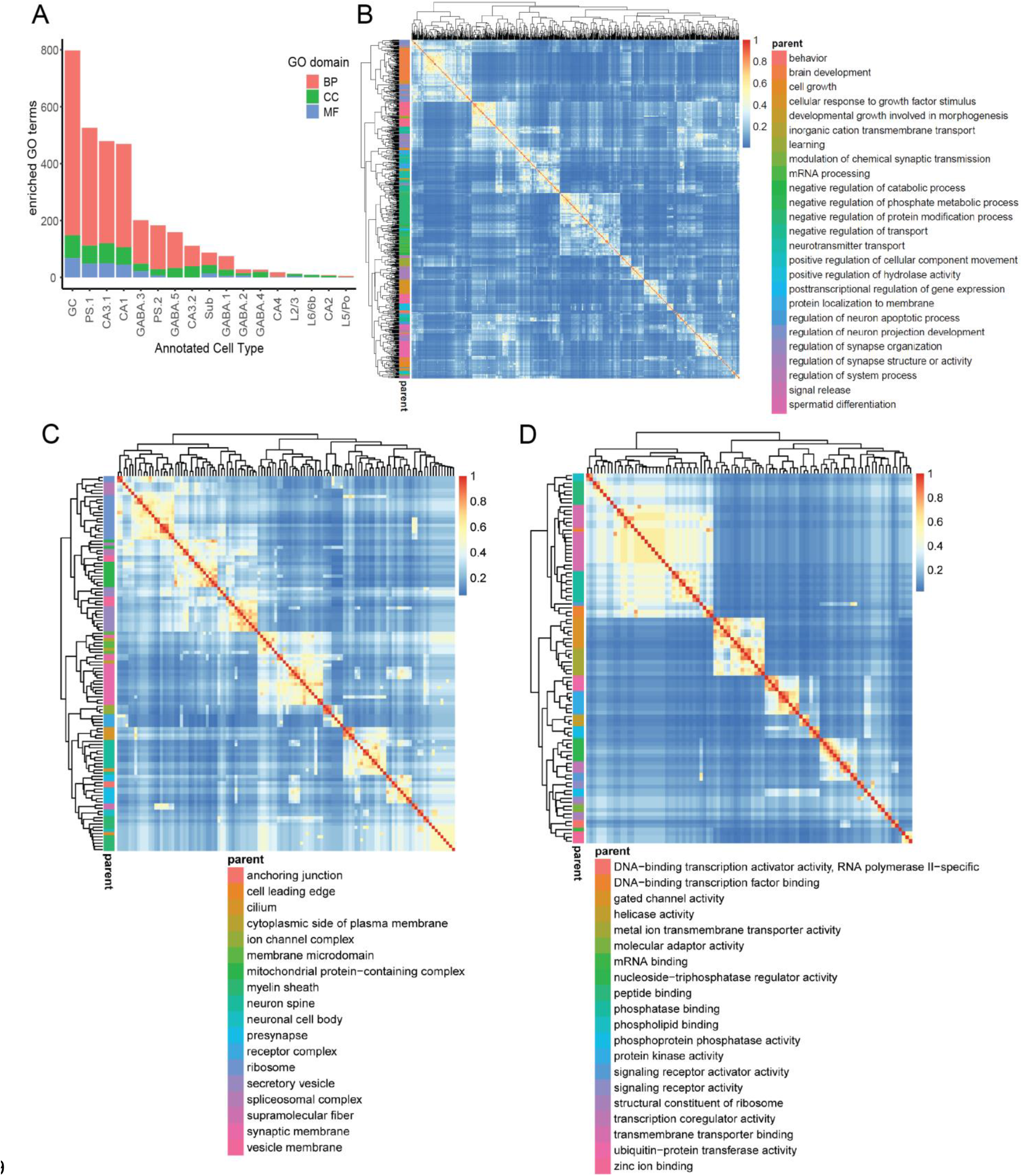
GO GSEA results and clustering. **(A)** Stacked bar plot illustrating number of GO terms from each domain (BP, CC, or MF) across all 17 cell types (“GC.1” and “GC.2” were combined into “GC” for purposes of GSEA). **(B)** Similarity matrix heatmap showing results from semantic similarity clustering analysis of all statistically significant BP GO terms, pooled across all cell types. **(C)** Same as B, but for all MF GO terms. **(D)** Same as B and C, but for all CC GO terms.

**Figure S6:**
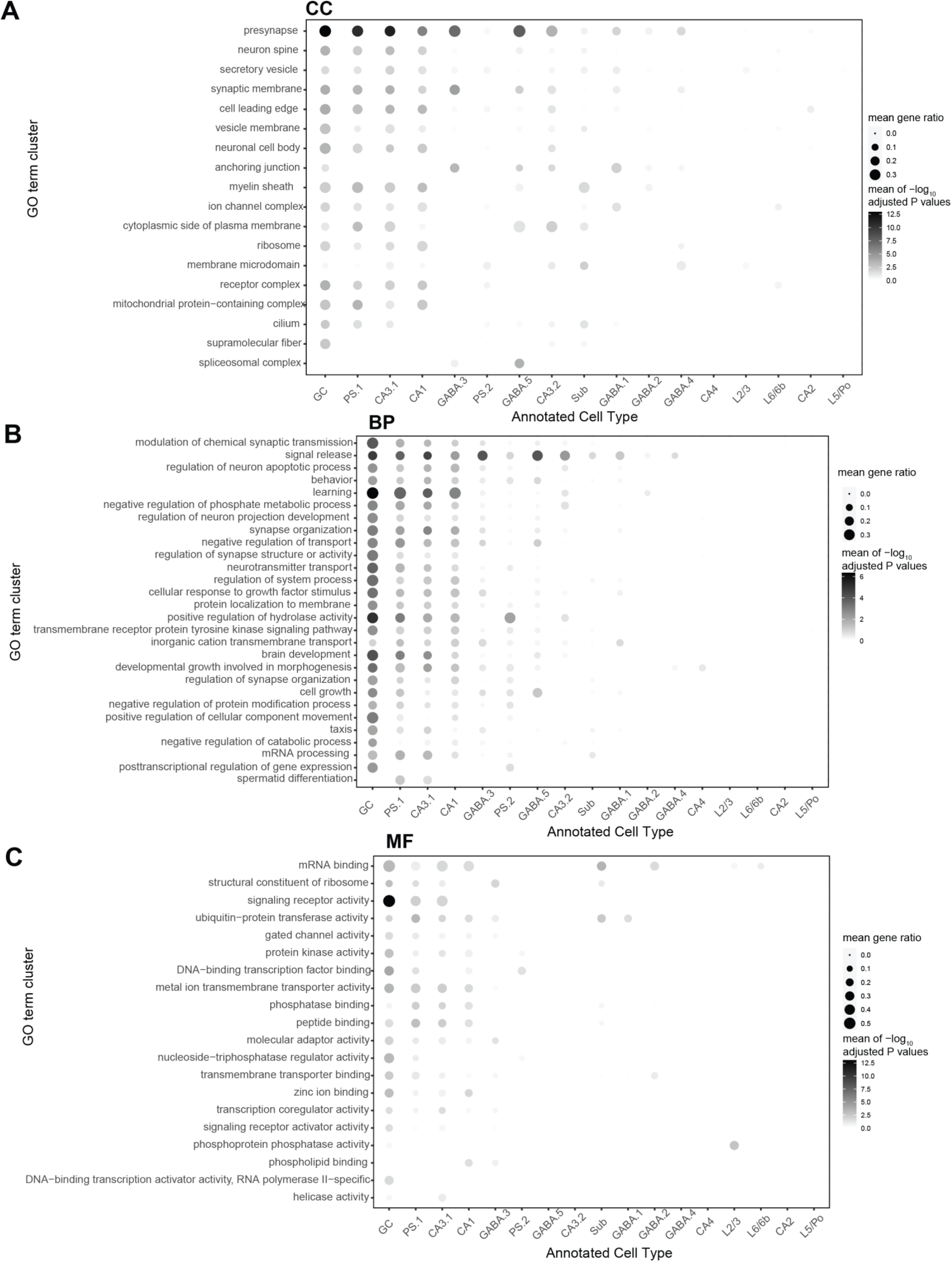
GO GSEA cluster overview. **(A)** Plot illustrating all Cellular Component (CC) GO term clusters obtained by semantic similarity analysis of cell type-specific GSEA results. Each row is a different CC GO cluster. Circles are shaded by the mean -log10 adjusted *p*-value, by cell type, for all terms within each cluster. Circles are sized by the mean gene ratio, by cell type, for all terms within each cluster. **(B)** Same as A, but for Biological Process (BP) GO term clusters. **(C)** Same as A&B, but for Molecular Function (MF) GO term clusters.

## Bibliography

Ahlmann-Eltze C. 2020. Combination Matrix Axis for “ggplot2” to Create “UpSet” Plots [R package ggupset version 0.3.0].

Alkadhi KA. 2019. Cellular and molecular differences between area CA1 and the dentate gyrus of the hippocampus. Mol Neurobiol 56:6566–6580.

Altar CA, Laeng P, Jurata LW, Brockman JA, Lemire A, Bullard J, Bukhman YV, Young TA, Charles V, Palfreyman MG. 2004. Electroconvulsive seizures regulate gene expression of distinct neurotrophic signaling pathways. J Neurosci 24:2667–2677.

Amezquita RA, Lun ATL, Becht E, Carey VJ, Carpp LN, Geistlinger L, Marini F, Rue-Albrecht K, Risso D, Soneson C, Waldron L, Pagès H, Smith ML, Huber W, Morgan M, Gottardo R, Hicks SC. 2020. Orchestrating single-cell analysis with Bioconductor. Nat Methods 17:137–145.

Andrade C, Singh NM, Thyagarajan S, Nagaraja N, Sanjay Kumar Rao N, Suresh Chandra J. 2008. Possible glutamatergic and lipid signalling mechanisms in ECT-induced retrograde amnesia: experimental evidence for involvement of COX-2, and review of literature. J Psychiatr Res 42:837–850.

Aoyama BB, Zanetti GG, Dias EV, Athié MCP, Lopes-Cendes I, Schwambach Vieira A. 2022. Transcriptomic analysis of dorsal and ventral subiculum after induction of acute seizures by electric stimulation of the perforant pathway in rats. Hippocampus 32:436–448.

Ashburner M, Ball CA, Blake JA, Botstein D, Butler H, Cherry JM, Davis AP, Dolinski K, Dwight SS, Eppig JT, Harris MA, Hill DP, Issel-Tarver L, Kasarskis A, Lewis S, Matese JC, Richardson JE, Ringwald M, Rubin GM, Sherlock G. 2000. Gene Ontology: tool for the unification of biology. Nat Genet 25:25–29.

Ayhan F, Kulkarni A, Berto S, Sivaprakasam K, Douglas C, Lega BC, Konopka G. 2021. Resolving cellular and molecular diversity along the hippocampal anterior-to-posterior axis in humans. Neuron 109:2091–2105.e6.

Bailey CH, Kandel ER, Harris KM. 2015. Structural components of synaptic plasticity and memory consolidation. Cold Spring Harb Perspect Biol 7:a021758.

Benjamini Y, Hochberg Y. 1995. Controlling the false discovery rate: a practical and powerful approach to multiple testing. Journal of the Royal Statistical Society: Series B (Methodological) 57:289–300.

Beurel E, Toups M, Nemeroff CB. 2020. The bidirectional relationship of depression and inflammation: double trouble. Neuron 107:234–256.

Bienkowski MS, Bowman I, Song MY, Gou L, Ard T, Cotter K, Zhu M, Benavidez NL, Yamashita S, Abu-Jaber J, Azam S, Lo D, Foster NN, Hintiryan H, Dong H-W. 2018. Integration of gene expression and brain-wide connectivity reveals the multiscale organization of mouse hippocampal networks. Nat Neurosci 21:1628–1643.

Böhm C, Peng Y, Maier N, Winterer J, Poulet JFA, Geiger JRP, Schmitz D. 2015. Functional Diversity of Subicular Principal Cells during Hippocampal Ripples. J Neurosci 35:13608–13618.

Bolwig TG. 2011. How does electroconvulsive therapy work? Theories on its mechanism. Can J Psychiatry 56:13–18.

Breheny P, Stromberg A, Lambert J. 2018. p-Value Histograms: Inference and Diagnostics. High-Throughput 7.

Carstens KE, Dudek SM. 2019. Regulation of synaptic plasticity in hippocampal area CA2. Curr Opin Neurobiol 54:194–199.

Castrén E, Monteggia LM. 2021. Brain-Derived Neurotrophic Factor Signaling in Depression and Antidepressant Action. Biol Psychiatry 90:128–136.

Cembrowski MS, Bachman JL, Wang L, Sugino K, Shields BC, Spruston N. 2016a. Spatial Gene-Expression Gradients Underlie Prominent Heterogeneity of CA1 Pyramidal Neurons. Neuron 89:351–368.

Cembrowski MS, Spruston N. 2019. Heterogeneity within classical cell types is the rule: lessons from hippocampal pyramidal neurons. Nat Rev Neurosci 20:193–204.

Cembrowski MS, Wang L, Lemire AL, Copeland M, DiLisio SF, Clements J, Spruston N. 2018. The subiculum is a patchwork of discrete subregions. eLife 7.

Cembrowski MS, Wang L, Sugino K, Shields BC, Spruston N. 2016b. Hipposeq: a comprehensive RNA-seq database of gene expression in hippocampal principal neurons. eLife 5:e14997.

Charlson F, Siskind D, Doi SAR, McCallum E, Broome A, Lie DC. 2012. ECT efficacy and treatment course: a systematic review and meta-analysis of twice vs thrice weekly schedules. J Affect Disord 138:1–8.

Chen Y, Lun ATL, Smyth GK. 2016. From reads to genes to pathways: differential expression analysis of RNA-Seq experiments using Rsubread and the edgeR quasi-likelihood pipeline. [version 2; peer review: 5 approved]. F1000Res 5:1438.

Chottekalapanda RU, Kalik S, Gresack J, Ayala A, Gao M, Wang W, Meller S, Aly A, Schaefer A, Greengard P. 2020. AP-1 controls the p11-dependent antidepressant response. Mol Psychiatry 25:1364–1381.

Collado-Torres L, Burke EE, Peterson A, Shin J, Straub RE, Rajpurohit A, Semick SA, Ulrich WS, BrainSeq Consortium, Price AJ, Valencia C, Tao R, Deep-Soboslay A, Hyde TM, Kleinman JE, Weinberger DR, Jaffe AE. 2019. Regional Heterogeneity in Gene Expression, Regulation, and Coherence in the Frontal Cortex and Hippocampus across Development and Schizophrenia. Neuron 103:203–216.e8.

Conway JR, Lex A, Gehlenborg N. 2017. UpSetR: an R package for the visualization of intersecting sets and their properties. Bioinformatics 33:2938–2940.

Cowley TR, Fahey B, O’Mara SM. 2008. COX-2, but not COX-1, activity is necessary for the induction of perforant path long-term potentiation and spatial learning in vivo. Eur J Neurosci 27:2999–3008.

Csardi G, Nepusz T. 2005. The Igraph Software Package for Complex Network Research. InterJournal Complex Systems 1695.

Dantzer R, O’Connor JC, Freund GG, Johnson RW, Kelley KW. 2008. From inflammation to sickness and depression: when the immune system subjugates the brain. Nat Rev Neurosci 9:46–56.

DeBruine ZJ, Melcher K, Triche TJ. 2021. Fast and robust non-negative matrix factorization for single-cell experiments. BioRxiv.

Dengler CG, Coulter DA. 2016. Normal and epilepsy-associated pathologic function of the dentate gyrus. Prog Brain Res 226:155–178.

Diering GH, Huganir RL. 2018. The AMPA receptor code of synaptic plasticity. Neuron 100:314–329.

Ding S-L, Yao Z, Hirokawa KE, Nguyen TN, Graybuck LT, Fong O, Bohn P, Ngo K, Smith KA, Koch C, Phillips JW, Lein ES, Harris JA, Tasic B, Zeng H. 2020. Distinct Transcriptomic Cell Types and Neural Circuits of the Subiculum and Prosubiculum along the Dorsal-Ventral Axis. Cell Rep 31:107648.

Dong H-W, Swanson LW, Chen L, Fanselow MS, Toga AW. 2009. Genomic-anatomic evidence for distinct functional domains in hippocampal field CA1. Proc Natl Acad Sci USA 106:11794–11799.

Dougherty KA, Islam T, Johnston D. 2012. Intrinsic excitability of CA1 pyramidal neurones from the rat dorsal and ventral hippocampus. J Physiol (Lond) 590:5707–5722.

Engin E, Zarnowska ED, Benke D, Tsvetkov E, Sigal M, Keist R, Bolshakov VY, Pearce RA, Rudolph U. 2015. Tonic Inhibitory Control of Dentate Gyrus Granule Cells by α5-Containing GABAA Receptors Reduces Memory Interference. J Neurosci 35:13698–13712.

Fertig EJ, Ding J, Favorov AV, Parmigiani G, Ochs MF. 2010. CoGAPS: an R/C++ package to identify patterns and biological process activity in transcriptomic data. Bioinformatics 26:2792–2793.

Freytag S, Lister R. 2020. schex avoids overplotting for large single-cell RNA-sequencing datasets. Bioinformatics 36:2291–2292.

Gallo FT, Katche C, Morici JF, Medina JH, Weisstaub NV. 2018. Immediate Early Genes, Memory and Psychiatric Disorders: Focus on c-Fos, Egr1 and Arc. Front Behav Neurosci 12:79.

Germain P-L, Lun A, Garcia Meixide C, Macnair W, Robinson MD. 2021. Doublet identification in single-cell sequencing data using scDblFinder. F1000Res 10:979.

Goldfarb S, Fainstein N, Ben-Hur T. 2020. Electroconvulsive stimulation attenuates chronic neuroinflammation. JCI Insight 5.

Graves AR, Moore SJ, Bloss EB, Mensh BD, Kath WL, Spruston N. 2012. Hippocampal pyramidal neurons comprise two distinct cell types that are countermodulated by metabotropic receptors. Neuron 76:776–789.

Gray JM, Spiegel I. 2019. Cell-type-specific programs for activity-regulated gene expression. Curr Opin Neurobiol 56:33–39.

Habib N, Avraham-Davidi I, Basu A, Burks T, Shekhar K, Hofree M, Choudhury SR, Aguet F, Gelfand E, Ardlie K, Weitz DA, Rozenblatt-Rosen O, Zhang F, Regev A. 2017. Massively parallel single-nucleus RNA-seq with DroNc-seq. Nat Methods 14:955–958.

Habib N, Li Y, Heidenreich M, Swiech L, Avraham-Davidi I, Trombetta JJ, Hession C, Zhang F, Regev A. 2016. Div-Seq: Single-nucleus RNA-Seq reveals dynamics of rare adult newborn neurons. Science 353:925–928.

Hainmueller T, Bartos M. 2020. Dentate gyrus circuits for encoding, retrieval and discrimination of episodic memories. Nat Rev Neurosci 21:153–168.

Harris KD, Hochgerner H, Skene NG, Magno L, Katona L, Bengtsson Gonzales C, Somogyi P, Kessaris N, Linnarsson S, Hjerling-Leffler J. 2018. Classes and continua of hippocampal CA1 inhibitory neurons revealed by single-cell transcriptomics. PLoS Biol 16:e2006387.

Hunt DL, Linaro D, Si B, Romani S, Spruston N. 2018. A novel pyramidal cell type promotes sharp-wave synchronization in the hippocampus. Nat Neurosci 21:985–995.

Hu P, Fabyanic E, Kwon DY, Tang S, Zhou Z, Wu H. 2017. Dissecting Cell-Type Composition and Activity-Dependent Transcriptional State in Mammalian Brains by Massively Parallel Single-Nucleus RNA-Seq. Mol Cell 68:1006–1015.e7.

Huber W, Carey VJ, Gentleman R, Anders S, Carlson M, Carvalho BS, Bravo HC, Davis S, Gatto L, Girke T, Gottardo R, Hahne F, Hansen KD, Irizarry RA, Lawrence M, Love MI, MacDonald J, Obenchain V, Oleś AK, Pagès H, Morgan M. 2015. Orchestrating high-throughput genomic analysis with Bioconductor. Nat Methods 12:115–121.

Imoto Y, Segi-Nishida E, Suzuki H, Kobayashi K. 2017. Rapid and stable changes in maturation-related phenotypes of the adult hippocampal neurons by electroconvulsive treatment. Mol Brain 10:8.

Jansson L, Wennström M, Johanson A, Tingström A. 2009. Glial cell activation in response to electroconvulsive seizures. Prog Neuropsychopharmacol Biol Psychiatry 33:1119–1128.

Jung MW, McNaughton BL. 1993. Spatial selectivity of unit activity in the hippocampal granular layer. Hippocampus 3:165–182.

Kalpachidou T, Makrygiannis AK, Pavlakis E, Stylianopoulou F, Chalepakis G, Stamatakis A. 2021. Behavioural effects of extracellular matrix protein Fras1 depletion in the mouse. Eur J Neurosci 53:3905–3919.

Kho KH, van Vreeswijk MF, Simpson S, Zwinderman AH. 2003. A meta-analysis of electroconvulsive therapy efficacy in depression. J ECT 19:139–147.

Kim Y, Spruston N. 2012. Target-specific output patterns are predicted by the distribution of regular-spiking and bursting pyramidal neurons in the subiculum. Hippocampus 22:693–706.

Kodama M, Russell DS, Duman RS. 2005. Electroconvulsive seizures increase the expression of MAP kinase phosphatases in limbic regions of rat brain. Neuropsychopharmacology 30:360–371.

Kotliar D, Veres A, Nagy MA, Tabrizi S, Hodis E, Melton DA, Sabeti PC. 2019. Identifying gene expression programs of cell-type identity and cellular activity with single-cell RNA-Seq. eLife 8.

Krieglstein K, Zheng F, Unsicker K, Alzheimer C. 2011. More than being protective: functional roles for TGF-β/activin signaling pathways at central synapses. Trends Neurosci 34:421–429.

Krueppel R, Remy S, Beck H. 2011. Dendritic integration in hippocampal dentate granule cells. Neuron 71:512–528.

Lacar B, Linker SB, Jaeger BN, Krishnaswami SR, Barron JJ, Kelder MJE, Parylak SL, Paquola ACM, Venepally P, Novotny M, O’Connor C, Fitzpatrick C, Erwin JA, Hsu JY, Husband D, McConnell MJ, Lasken R, Gage FH. 2016. Nuclear RNA-seq of single neurons reveals molecular signatures of activation. Nat Commun 7:11022.

Lee H, Wang C, Deshmukh SS, Knierim JJ. 2015. Neural Population Evidence of Functional Heterogeneity along the CA3 Transverse Axis: Pattern Completion versus Pattern Separation. Neuron 87:1093–1105.

Lee M, Yoon J, Song H, Lee B, Lam DT, Yoon J, Baek K, Clevers H, Jeong Y. 2017. Tcf7l2 plays crucial roles in forebrain development through regulation of thalamic and habenular neuron identity and connectivity. Dev Biol 424:62–76.

Leutgeb JK, Leutgeb S, Moser M-B, Moser EI. 2007. Pattern separation in the dentate gyrus and CA3 of the hippocampus. Science 315:961–966.

Li ZX, Yu HM, Jiang KW. 2013. Tonic GABA inhibition in hippocampal dentate granule cells: its regulation and function in temporal lobe epilepsies. Acta Physiol (Oxf) 209:199–211.

López DE, Ballaz SJ. 2020. The Role of Brain Cyclooxygenase-2 (Cox-2) Beyond Neuroinflammation: Neuronal Homeostasis in Memory and Anxiety. Mol Neurobiol 57:5167–5176.

Lund SP, Nettleton D, McCarthy DJ, Smyth GK. 2012. Detecting differential expression in RNA-sequence data using quasi-likelihood with shrunken dispersion estimates. Stat Appl Genet Mol Biol 11.

Lun A. 2022. bluster: Clustering Algorithms for Bioconductor. R package version 1.8.0. Bioconductor.

Lun ATL, Bach K, Marioni JC. 2016a. Pooling across cells to normalize single-cell RNA sequencing data with many zero counts. Genome Biol 17:75.

Lun ATL, Chen Y, Smyth GK. 2016b. It’s DE-licious: A Recipe for Differential Expression Analyses of RNA-seq Experiments Using Quasi-Likelihood Methods in edgeR. Methods Mol Biol 1418:391–416.

Lun ATL, Riesenfeld S, Andrews T, Dao TP, Gomes T, participants in the 1st Human Cell Atlas Jamboree, Marioni JC. 2019. EmptyDrops: distinguishing cells from empty droplets in droplet-based single-cell RNA sequencing data. Genome Biol 20:63.

Marín-Burgin A, Mongiat LA, Pardi MB, Schinder AF. 2012. Unique processing during a period of high excitation/inhibition balance in adult-born neurons. Science 335:1238–1242.

Maynard KR, Hobbs JW, Rajpurohit SK, Martinowich K. 2018. Electroconvulsive seizures influence dendritic spine morphology and BDNF expression in a neuroendocrine model of depression. Brain Stimulat 11:856–859.

McCarthy DJ, Campbell KR, Lun ATL, Wills QF. 2017. Scater: pre-processing, quality control, normalization and visualization of single-cell RNA-seq data in R. Bioinformatics 33:1179–1186.

McCarthy DJ, Chen Y, Smyth GK. 2012. Differential expression analysis of multifactor RNA-Seq experiments with respect to biological variation. Nucleic Acids Res 40:4288–4297.

McInnes L, Healy J, Melville J. 2018. UMAP: Uniform Manifold Approximation and Projection for Dimension Reduction. arXiv.

Meyers EA, Kessler JA. 2017. TGF-β Family Signaling in Neural and Neuronal Differentiation, Development, and Function. Cold Spring Harb Perspect Biol 9.

Miller AH, Raison CL. 2016. The role of inflammation in depression: from evolutionary imperative to modern treatment target. Nat Rev Immunol 16:22–34.

Mitra S, Werner C, Dietz DM. 2022. Neuroadaptations and TGF-β signaling: emerging role in models of neuropsychiatric disorders. Mol Psychiatry 27:296–306.

Nagalski A, Puelles L, Dabrowski M, Wegierski T, Kuznicki J, Wisniewska MB. 2016. Molecular anatomy of the thalamic complex and the underlying transcription factors. Brain Struct Funct 221:2493–2510.

Nagy C, Vaillancourt K, Turecki G. 2018. A role for activity-dependent epigenetics in the development and treatment of major depressive disorder. Genes Brain Behav 17:e12446.

Newton SS, Collier EF, Hunsberger J, Adams D, Terwilliger R, Selvanayagam E, Duman RS. 2003. Gene profile of electroconvulsive seizures: induction of neurotrophic and angiogenic factors. J Neurosci 23:10841–10851.

Ongür D, Pohlman J, Dow AL, Eisch AJ, Edwin F, Heckers S, Cohen BM, Patel TB, Carlezon WA. 2007. Electroconvulsive seizures stimulate glial proliferation and reduce expression of Sprouty2 within the prefrontal cortex of rats. Biol Psychiatry 62:505–512.

Palacios-Filardo J, Mellor JR. 2019. Neuromodulation of hippocampal long-term synaptic plasticity. Curr Opin Neurobiol 54:37–43.

Pardi MB, Ogando MB, Schinder AF, Marin-Burgin A. 2015. Differential inhibition onto developing and mature granule cells generates high-frequency filters with variable gain. eLife 4:e08764.

Pelkey KA, Chittajallu R, Craig MT, Tricoire L, Wester JC, McBain CJ. 2017. Hippocampal gabaergic inhibitory interneurons. Physiol Rev 97:1619–1747.

Ploski JE, Newton SS, Duman RS. 2006. Electroconvulsive seizure-induced gene expression profile of the hippocampus dentate gyrus granule cell layer. J Neurochem 99:1122–1132.

Robinson MD, McCarthy DJ, Smyth GK. 2010. edgeR: a Bioconductor package for differential expression analysis of digital gene expression data. Bioinformatics 26:139–140.

Rosenthal EH, Tonchev AB, Stoykova A, Chowdhury K. 2012. Regulation of archicortical arealization by the transcription factor Zbtb20. Hippocampus 22:2144–2156.

Rue-Albrecht K, Marini F, Soneson C, Lun ATL. 2018. iSEE: Interactive SummarizedExperiment Explorer. [version 1; peer review: 3 approved]. F1000Res 7:741.

Sayols S. 2020. rrvgo: a Bioconductor package to reduce and visualize Gene Ontology terms. Bioconductor.

Schloesser RJ, Orvoen S, Jimenez DV, Hardy NF, Maynard KR, Sukumar M, Manji HK, Gardier AM, David DJ, Martinowich K. 2015. Antidepressant-like Effects of Electroconvulsive Seizures Require Adult Neurogenesis in a Neuroendocrine Model of Depression. Brain Stimulat 8:862–867.

Sherman TD, Gao T, Fertig EJ. 2020. CoGAPS 3: Bayesian non-negative matrix factorization for single-cell analysis with asynchronous updates and sparse data structures. BMC Bioinformatics 21:453.

Stark DT, Bazan NG. 2011. Synaptic and extrasynaptic NMDA receptors differentially modulate neuronal cyclooxygenase-2 function, lipid peroxidation, and neuroprotection. J Neurosci 31:13710–13721.

Stein-O’Brien GL, Arora R, Culhane AC, Favorov AV, Garmire LX, Greene CS, Goff LA, Li Y, Ngom A, Ochs MF, Xu Y, Fertig EJ. 2018. Enter the Matrix: Factorization Uncovers Knowledge from Omics. Trends Genet 34:790–805.

Stein-O’Brien GL, Carey JL, Lee WS, Considine M, Favorov AV, Flam E, Guo T, Li S, Marchionni L, Sherman T, Sivy S, Gaykalova DA, McKay RD, Ochs MF, Colantuoni C, Fertig EJ. 2017. PatternMarkers & GWCoGAPS for novel data-driven biomarkers via whole transcriptome NMF. Bioinformatics 33:1892–1894.

Stein-O’Brien GL, Clark BS, Sherman T, Zibetti C, Hu Q, Sealfon R, Liu S, Qian J, Colantuoni C, Blackshaw S, Goff LA, Fertig EJ. 2019. Decomposing Cell Identity for Transfer Learning across Cellular Measurements, Platforms, Tissues, and Species. Cell Syst 8:395–411.e8.

Steward O. 1994. Electroconvulsive seizures upregulate astroglial gene expression selectively in the dentate gyrus. Brain Res Mol Brain Res 25:217–224.

Subramanian A, Tamayo P, Mootha VK, Mukherjee S, Ebert BL, Gillette MA, Paulovich A, Pomeroy SL, Golub TR, Lander ES, Mesirov JP. 2005. Gene set enrichment analysis: a knowledge-based approach for interpreting genome-wide expression profiles. Proc Natl Acad Sci USA 102:15545–15550.

Sun Q, Sotayo A, Cazzulino AS, Snyder AM, Denny CA, Siegelbaum SA. 2017. Proximodistal Heterogeneity of Hippocampal CA3 Pyramidal Neuron Intrinsic Properties, Connectivity, and Reactivation during Memory Recall. Neuron 95:656–672.e3.

The Gene Ontology Consortium. 2021. The Gene Ontology resource: enriching a GOld mine. Nucleic Acids Res 49:D325–D334.

Thompson CL, Pathak SD, Jeromin A, Ng LL, MacPherson CR, Mortrud MT, Cusick A, Riley ZL, Sunkin SM, Bernard A, Puchalski RB, Gage FH, Jones AR, Bajic VB, Hawrylycz MJ, Lein ES. 2008. Genomic anatomy of the hippocampus. Neuron 60:1010–1021.

Townes FW, Hicks SC, Aryee MJ, Irizarry RA. 2019. Feature selection and dimension reduction for single-cell RNA-Seq based on a multinomial model. Genome Biol 20:295.

Tran MN, Maynard KR, Spangler A, Huuki LA, Montgomery KD, Sadashivaiah V, Tippani M, Barry BK, Hancock DB, Hicks SC, Kleinman JE, Hyde TM, Collado-Torres L, Jaffe AE, Martinowich K. 2021. Single-nucleus transcriptome analysis reveals cell-type-specific molecular signatures across reward circuitry in the human brain. Neuron 109:3088–3103.e5.

Tyssowski KM, DeStefino NR, Cho J-H, Dunn CJ, Poston RG, Carty CE, Jones RD, Chang SM, Romeo P, Wurzelmann MK, Ward JM, Andermann ML, Saha RN, Dudek SM, Gray JM. 2018. Different neuronal activity patterns induce different gene expression programs. Neuron 98:530–546.e11.

Tyssowski KM, Gray JM. 2019. The neuronal stimulation-transcription coupling map. Curr Opin Neurobiol 59:87–94.

Ueno M, Sugimoto M, Ohtsubo K, Sakai N, Endo A, Shikano K, Imoto Y, Segi-Nishida E. 2019. The effect of electroconvulsive seizure on survival, neuronal differentiation, and expression of the maturation marker in the adult mouse hippocampus. J Neurochem 149:488–498.

UK ECT Review Group. 2003. Efficacy and safety of electroconvulsive therapy in depressive disorders: a systematic review and meta-analysis. Lancet 361:799–808.

Wang JZ, Du Z, Payattakool R, Yu PS, Chen C-F. 2007. A new method to measure the semantic similarity of GO terms. Bioinformatics 23:1274–1281.

Wennström M, Hellsten J, Ekdahl CT, Tingström A. 2003. Electroconvulsive seizures induce proliferation of NG2-expressing glial cells in adult rat hippocampus. Biol Psychiatry 54:1015–1024.

Wu T, Hu E, Xu S, Chen M, Guo P, Dai Z, Feng T, Zhou L, Tang W, Zhan L, Fu X, Liu S, Bo X, Yu G. 2021. clusterProfiler 4.0: A universal enrichment tool for interpreting omics data. Innovation (Camb) 2:100141.

Wu YE, Pan L, Zuo Y, Li X, Hong W. 2017. Detecting Activated Cell Populations Using Single-Cell RNA-Seq. Neuron 96:313–329.e6.

Xu C, Su Z. 2015. Identification of cell types from single-cell transcriptomes using a novel clustering method. Bioinformatics 31:1974–1980.

Yao Z, van Velthoven CTJ, Nguyen TN, Goldy J, Sedeno-Cortes AE, Baftizadeh F, Bertagnolli D, Casper T, Chiang M, Crichton K, Ding S-L, Fong O, Garren E, Glandon A, Gouwens NW, Gray J, Graybuck LT, Hawrylycz MJ, Hirschstein D, Kroll M, Zeng H. 2021. A taxonomy of transcriptomic cell types across the isocortex and hippocampal formation. Cell 184:3222–3241.e26.

Yu G, Li F, Qin Y, Bo X, Wu Y, Wang S. 2010. GOSemSim: an R package for measuring semantic similarity among GO terms and gene products. Bioinformatics 26:976–978.

Yu G, Wang L-G, Han Y, He Q-Y. 2012. clusterProfiler: an R package for comparing biological themes among gene clusters. OMICS 16:284–287.

Yu G. 2020. Gene ontology semantic similarity analysis using gosemsim. Methods Mol Biol 2117:207–215.

Zeisel A, Muñoz-Manchado AB, Codeluppi S, Lönnerberg P, La Manno G, Juréus A, Marques S, Munguba H, He L, Betsholtz C, Rolny C, Castelo-Branco G, Hjerling-Leffler J, Linnarsson S. 2015. Brain structure. Cell types in the mouse cortex and hippocampus revealed by single-cell RNA-seq. Science 347:1138–1142.

Zheng GXY, Terry JM, Belgrader P, Ryvkin P, Bent ZW, Wilson R, Ziraldo SB, Wheeler TD, McDermott GP, Zhu J, Gregory MT, Shuga J, Montesclaros L, Underwood JG, Masquelier DA, Nishimura SY, Schnall-Levin M, Wyatt PW, Hindson CM, Bharadwaj R, Bielas JH. 2017. Massively parallel digital transcriptional profiling of single cells. Nat Commun 8:14049.

